# Functional Locality–Aligned Learning Reveals Structure–Function Causality in Enzyme Kinetics

**DOI:** 10.64898/2026.03.04.709726

**Authors:** Hao Zhang, He Zhang, Miao Kang, Kaipeng Zhang, Tao Yang, Nanning Zheng

## Abstract

Accurate estimation of enzyme kinetic parameters is essential for enzyme engineering and industrial biocatalysis, yet their experimental measurement remains labor-intensive and costly. Although machine learning offers an efficient alternative, existing methods still struggle to generalize to unseen enzymes and substrates. In particular, current three-dimensional (3D) structure–aware approaches rely on whole-enzyme 3D geometric structures and neglect substrate 3D geometry, often yielding limited or even degraded performance. We identify a fundamental limitation underlying these methods: a mismatch between the structural representation learning scale and the functional locality scale, which weakens structure–function causality in enzyme kinetics. To address this issue, we introduce *EnzymePlex*, a functional locality–aligned framework that aligns inductive biases with the localized structural determinants of enzyme function by prioritizing catalytic pockets, integrating substrate 3D geometry, and modeling nuanced enzyme–substrate interplay under the guidance of pocket-level structural priors. EnzymePlex achieves state-of-the-art performance across multiple benchmarks and substantially improves generalization under stringent out-of-distribution evaluations. Beyond predictive accuracy, EnzymePlex learns mechanistically aligned representations, with attention enriched at catalytic pocket residues and substrate reaction centers despite receiving no explicit supervision for either. Moreover, when applied to recently reported wet-lab data, EnzymePlex effectively prioritizes high-activity enzyme variants and identifies potent inhibitors, highlighting its potential to accelerate enzyme engineering and drug discovery.

## 1 Main

Enzymes are central biocatalysts that sustain diverse biological processes and underpin a wide range of biotechnological applications, including pharmaceutical manufacturing [1, 2], food and beverage bioprocessing [3, 4], and renewable biofuel production [5, 6]. Quantitative characterization of enzyme kinetics, most prominently turnover numbers (k_cat_), Michaelis constants (K_m_), and inhibition constants (K_i_), is essential for understanding catalytic efficiency and substrate affinity, and for facilitating downstream tasks such as enzyme engineering and drug discovery [7–10]. However, experimental determination of these enzyme kinetic parameters relies on costly and labor-intensive *in vitro* assays, posing a significant bottleneck for high-throughput enzyme optimization and directed evolution.

Advances in machine learning (ML) have profoundly transformed biomolecular research [11–14], offering an efficient alternative for predicting enzyme kinetic parameters directly from enzyme–substrate pairs. Early work by Heckmann et al. [15] employed ML models with handcrafted biochemical features to predict k_cat_ of enzymes from *Escherichia coli*. Later, Li et al. [16] introduced DLKcat, which predicted k_cat_ across diverse organisms by encoding enzyme sequences and substrate two-dimensional (2D) molecular graphs. The emergence of large protein and molecular language models [17–21] further enhanced prediction performance by leveraging pretrained representations of enzyme sequences and substrate SMILES [8, 22, 23].

However, these coarse-grained representations lack 3D geometric structural information for both enzymes and substrates, which fundamentally determines enzyme catalytic function [8, 24, 25]. This limitation can constrain their generalization to enzymes and substrates previously unseen during training. Recent studies have sought to incorporate enzyme 3D structural information into ML models, yet empirical evidence shows that using whole-enzyme structures often yields only marginal gains and can even degrade performance. For instance, CatPred [26] used an SE(3)-equivariant graph neural network [27] to encode whole-enzyme structures but achieved only marginal performance improvement for K_i_ and a decrease in k_cat_. CatPro [8] leveraged 3D structure–informed pretrained models [28] to derive whole-enzyme structural embeddings but observed a slight reduction in k_cat_ accuracy and negligible gains for K_m_.

We posit that this limitation reflects a fundamental mismatch between the scale at which representations are learned and the functional locality scale, *i.e.*, the localized structural determinants of catalytic function. Specifically, enzyme kinetics are strongly shaped by localized enzyme–substrate interactions within the catalytic pocket environment [29, 30], so when representation learning is uniformly applied to the entire enzyme structure, the pocket-level features that underlie structure–function causality can be diluted by functionally irrelevant or weakly related regions. To address this challenge, we introduce *EnzymePlex*, a functional locality–aligned framework that explicitly conditions representation learning on catalytic pocket environments. EnzymePlex employs a dedicated SE(3)-invariant graph encoder to extract structural priors from both localized pocket structures and pocket surfaces, the latter providing a concise and functionally meaningful description of enzymes as continuous molecular shapes enriched with geometric and chemical properties [31]. In parallel, substrates are encoded using a 3D molecular structure encoder pretrained on large-scale molecular conformation datasets [32], providing structure-informed inductive biases by explicitly modeling substrate geometry. Finally, EnzymePlex integrates enzyme–pocket, substrate, and enzyme–sequence features through a multi-modal interaction module guided by pocket-level structural priors, producing catalysis-aware representations for enzyme kinetic parameter prediction.

EnzymePlex achieves state-of-the-art (SOTA) performance across standard benchmarks for k_cat_, K_m_, and K_i_ prediction and demonstrates strong out-of-distribution generalization to both enzymes and substrates with low similarity to those in training. Under a 40% sequence-identity cutoff, EnzymePlex improves the coefficient of determination (*R*^2^) over the second-best method by 5.71%, 3.00%, and 15.60% for k_cat_, K_m_, and K_i_, respectively; under a 40% molecular similarity cutoff, these gains rise to 11.07%, 36.25%, and 33.42%. Beyond predictive accuracy, EnzymePlex further learns mechanistically aligned representations, with attention enriched at known catalytic pocket residues and molecular reaction centers, despite receiving no explicit supervision for either. Moreover, EnzymePlex demonstrates practical utility on recently reported wet-lab measurements, successfully prioritizing high-activity enzyme variants and identifying potent inhibitors. Collectively, these findings underscore that functional locality–aligned representation learning offers a principled and effective computational framework for robust prediction in structure-aware biomolecular systems, and provide a powerful tool for enzyme engineering and drug discovery.

## 2 Results

### 2.1 Workflow of EnzymePlex

EnzymePlex takes enzyme 3D structures and sequences, together with substrate 3D structures, as inputs to predict enzyme kinetic parameters (Fig. 1a). To test whether aligning representation learning with functional locality improves robustness and generalization, the framework is designed to condition multi-modal interaction modeling on catalytically localized structural priors. Specifically, we first identify the enzymes’ binding pockets and then derive their corresponding pocket structures (Sec. 4.4.1). We then augment these structures with surface-based geometric and physicochemical descriptors, including shape index, distance-dependent curvature, hydrophobicity, hydrogen-bonding potential, and Poisson–Boltzmann electrostatics(Fig. 1b; Sec. 4.4.1), as pocket surfaces constitute the direct interface for substrate binding and encode informative features for fine-grained biomolecular interactions [31, 33]. For substrates, initial conformations are generated using RDKit [34] and encoded with Uni-Mol2-570M [32], a large-scale pretrained 3D molecular structure model that provides geometry-informed substrate representations. In parallel, residue-level enzyme sequence representations are extracted using a pretrained protein language model, consistent with prior approaches [26].

**Fig. 1:**
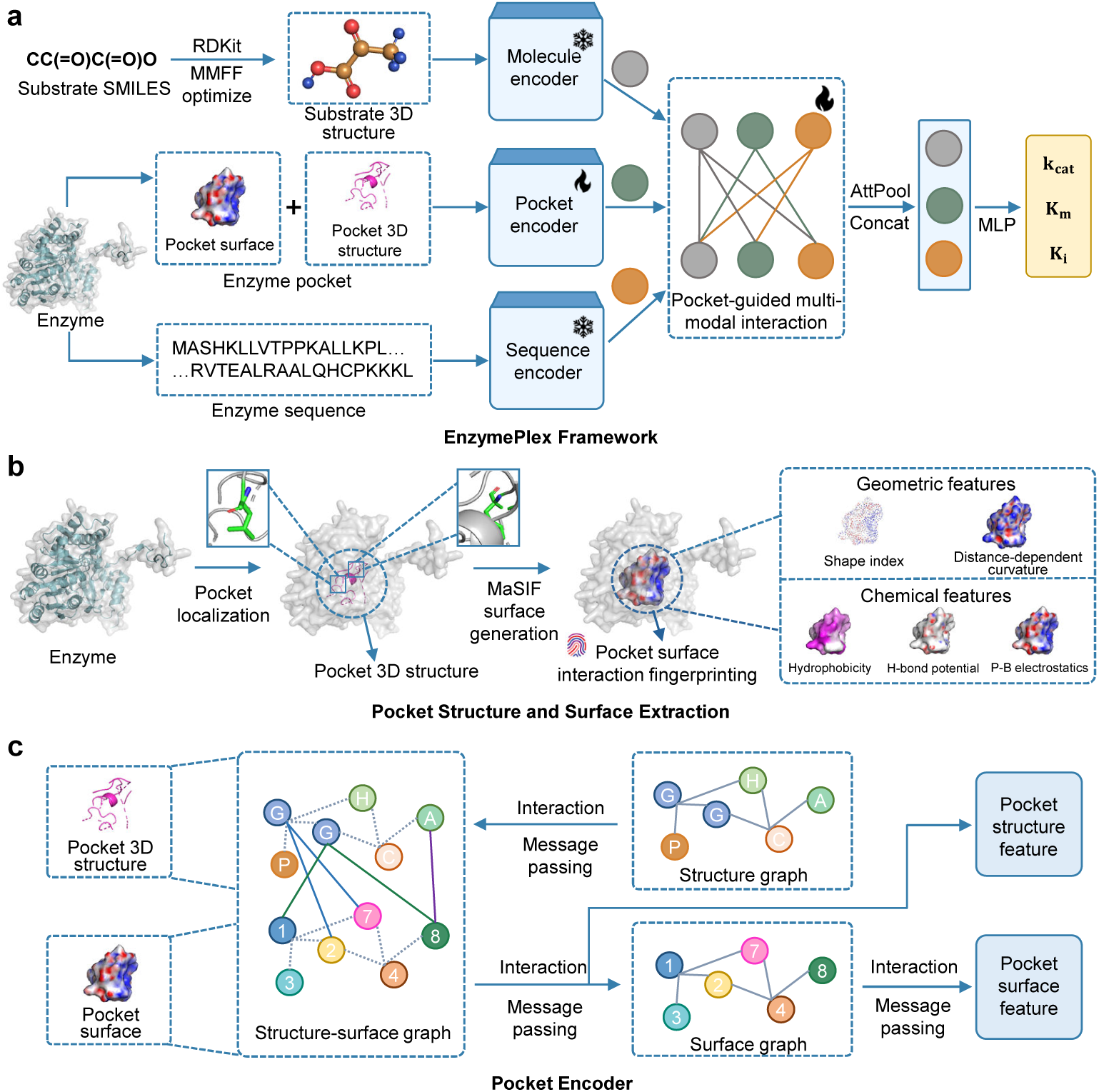
Overview of EnzymePlex. **(a)** EnzymePlex integrates a pretrained 3D molecular structure encoder, an SE(3)-invariant pocket encoder, and a pretrained protein sequence encoder to derive substrate-, pocket-, and sequence-level representations. Trainable modules are shown with flame icons, frozen ones with snowflakes. Substrate 3D conformations are generated using RDKit and subsequently refined with the MMFF94 force field. A pocket-guided multi-modal interaction module captures the biochemical interaction patterns linking the enzyme pocket, substrate molecule, and enzyme sequence. Attention pooling (AttPool), followed by a multilayer perceptron (MLP), produces the predicted kinetic parameters k_cat_, K_m_, and K_i_. **(b)** Catalytic pockets are first identified, and the resulting 3D pocket structures are subsequently used to compute MaSIF-based surface geometric and physicochemical properties. **(c)** An SE(3)-invariant graph neural network encoder models the enzyme pocket by performing message passing over the pocket structure graph, the structure–surface interaction graph, and the pocket surface graph.

Whereas many existing enzyme kinetics predictors rely on simple feature fusion strategies, such as direct concatenation [8, 26], limiting their ability to capture catalytically relevant interaction patterns. To address this limitation, EnzymePlex employs a multi-modal interaction module that integrates enzyme and substrate representations under the guidance of pocket-level functional locality (Fig. 1c; see Sec. 4.4.5). The resulting representations are aggregated via attention pooling, concatenated, and passed to a multilayer perceptron (MLP) to predict the respective targets (adopting log10-transformed kinetic parameters following prior works [26]). Unless otherwise specified, we trained EnzymePlex and all baseline models on CatPred-DB [26], an enzyme kinetics dataset curated from BRENDA [35], SABIO-RK [36], and UniProt [37]. More data processing and model implementation details are provided in Sec. 4.1 and Sec. 4.4.

### 2.2 Performance on enzyme kinetic parameter prediction

We first evaluated EnzymePlex on enzyme–substrate pairs in the CatPred-DB test set that were unseen during training. We compared EnzymePlex with DLKcat [16], UniKP [22], CataPro [8], and CatPred [26]. The former three represent SOTA sequence- and SMILES-based representation learning approaches, whereas the latter incorporates global structural representations derived from whole-enzyme structures. Performance was assessed using the coefficient of determination (*R*^2^) and mean absolute error (MAE) relative to experimental measurements (see Sec. 4.3). As shown in Fig. 2a and Supplementary Fig. S1a, EnzymePlex consistently outperformed all baselines on test samples across the three kinetic parameters. On k_cat_, K_m_, and K_i_ test sets, EnzymePlex achieved *R*^2^ values of 0.634, 0.671, and 0.680, and MAE values of 0.657, 0.520, and 0.783, respectively, corresponding to relative improvements of 3.43%, 3.39%, and 6.75% in *R*^2^, and reductions of 5.74%, 5.63%, and 9.48% in MAE compared with the second-best method. Ablation analyses indicate that the pocket-guided multi-modal interaction module contributes substantially to performance gains (Fig. 2b and Supplementary Fig. S1b), particularly under more stringent out-of-distribution (OOD) settings (see more results in Sec. 2.3). In contrast, CatPred exhibited limited or even degraded performance when incorporating structural features from entire enzyme 3D structures [26], underscoring the importance of conditioning representation learning on catalytically localized structures when functional determinants are spatially sparse. Next, we examined whether EnzymePlex captured established biochemical patterns. For metabolic trends, enzymes involved in primary central and energy metabolism exhibited significantly higher predicted k_cat_ values (median 14.494 s^−1^) than those associated with intermediary and secondary metabolism (median 11.926 s^−1^; *P* = 2.4 × 10^−4^; Fig. 2c), consistent with observations that central metabolic enzymes evolve higher turnover to sustain essential fluxes [39]. Primary metabolism enzymes also displayed higher predicted K_m_ values (median 0.256 mM vs. 0.124 mM; *P* = 1.3 × 10 ; Supplementary Fig. S1c) and higher predicted K_i_ values (median 0.084 mM vs. 0.036 mM; *P* = 1.7 10^−14^; Supplementary Fig. S1d), consistent with experimental annotations and indicative of comparatively weaker substrate and inhibitor binding. To assess enzyme promiscuity, we grouped substrates into preferred and alternative categories based on experimental annotations. EnzymePlex predicted significantly higher k_cat_ values for preferred substrates (median 15.227 s^−1^; *P* = 1.6×10 ; Supplementary Fig. S1e) and correspondingly lower K_m_ values (median 0.084 mM; *P* = 6.3×10 ; Fig. 2d), reflecting enhanced catalytic efficiency and tighter binding. Finally, for substrate-associated enzyme diversity, enzymes were categorized as native or underground based on experimental K_i_ measurements. EnzymePlex predicted lower K_i_ values for native enzymes (median 0.031 mM; *P* = 1.6 10^−4^; Fig. 2e), indicating that the model captures expected differences in inhibitory sensitivity across enzyme groups.

**Fig. 2:**
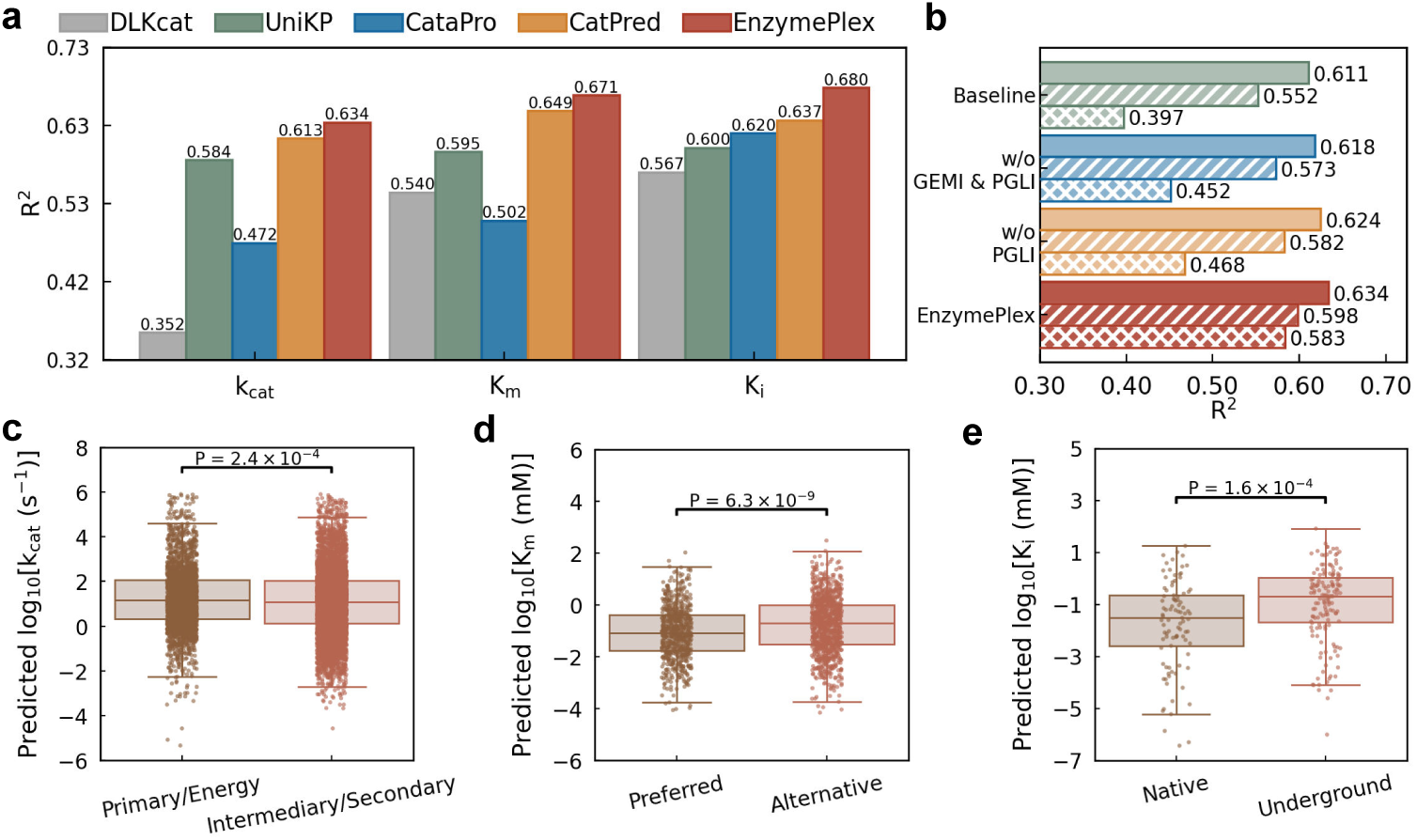
Performance of EnzymePlex. **(a)** Benchmark comparison of EnzymePlex with other methods on the CatPred-DB test set. Coefficient of determination (*R*^2^) values are separately reported for k_cat_, K_m_, and K_i_. **(b)** Performance of EnzymePlex variants for k_cat_ prediction on the test set (solid bars) and molecule-level OOD subsets. Two molecule-level OOD subsets were defined using molecular similarity cutoffs of 99% (diagonal-hatched bars) and 40% (cross-hatched bars), respectively. The Baseline variant excludes both the pocket-guided multi-modal interaction module and the 3D molecular structure encoder, and instead adopts the 2D molecular graph encoder [38] used in CatPred. GEMI: global enzyme-molecule interaction; PGLI: pocket-prior guided local interaction. **(c)** Comparison of EnzymePlex performance on k_cat_ for enzymes involved in primary central and energy metabolism versus intermediary and secondary metabolism. **(d)** Enzyme promiscuity analysis for K_m_. For enzymes acting on multiple substrates, substrates were grouped into preferred and alternative categories based on experimentally measured kinetics. **(e)** Substrate-associated enzyme diversity analysis for K_i_. For substrates linked to multiple enzymes, enzymes were classified as native or underground according to their inhibition constants. Two-sided Wilcoxon rank-sum tests were used to compute *P* values. In each box plot, the central line denotes the median, the box shows the interquartile range, and whiskers extend to 1.5 times the interquartile range.

### 2.3 Functional locality-aligned learning leads to improved out-of-distribution generalization

To evaluate the robustness of EnzymePlex under distribution shift, we assessed its performance on progressively stringent enzyme- and molecule-level OOD settings. Both OOD splits were constructed using enzyme sequence identity and molecular structural similarity thresholds of 99%, 80%, 60%, and 40%, where lower thresholds correspond to increasingly challenging generalization scenarios (see Sec. 4.1).

For enzyme-level OOD generalization, EnzymePlex consistently achieved higher *R*^2^ values than all baselines across sequence identity cutoffs and kinetic parameters (Fig. 3a), demonstrating robust performance on dissimilar enzyme sequences. On the K_i_ test set in particular, EnzymePlex surpassed the strongest competing method by 19.07%, 25.33%, 34.35%, and 15.60% across the four similarity thresholds, respectively. A similar pattern was observed for molecule-level OOD generalization (Fig. 3b).

**Fig. 3:**
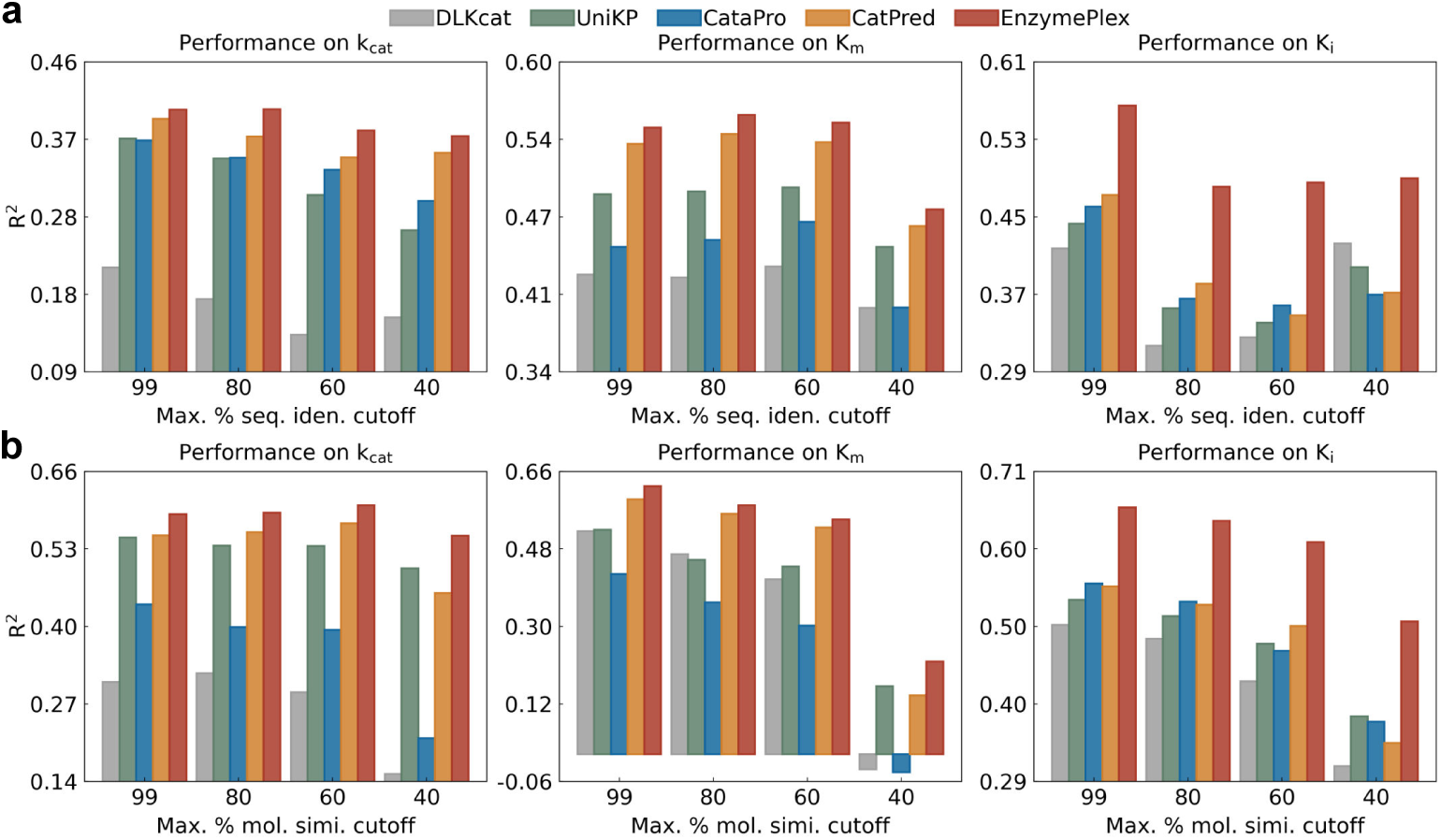
Out-of-distribution performance of EnzymePlex compared with other methods. *R*^2^ values are reported as a function of decreasing enzyme sequence identity or substrate molecular similarity cutoffs. OOD subsets are constructed from the CatPred test set by excluding samples whose similarity to the training set exceeds the specified threshold. **(a)** Enzyme-level OOD performance under sequence identity cutoffs. **(b)** Molecule-level OOD performance under molecular similarity cutoffs.

EnzymePlex consistently outperformed baseline models across all molecular similarity cutoffs and kinetic parameters. For K_i_ prediction in particular, EnzymePlex exceeded the strongest competing method by 18.28%, 20.22%, 22.36%, and 33.42% across progressively stricter similarity thresholds. Notably, as molecular similarity to the training distribution decreased, the performance gap between the global structural approach represented by CatPred (for K_i_) and the locality-aligned framework widens substantially, indicating improved robustness of EnzymePlex under increasingly severe distribution shifts. Consistent trends were also observed in terms of the MAE metric (see Supplementary A.1).

Collectively, these findings highlight the intrinsic difficulty of enzyme- and molecule-level OOD generalization for methods relying on global structural representations. By prioritizing pocket-level structural priors and incorporating substrate 3D geometries, EnzymePlex significantly mitigates this challenge(Fig. 2b and Supplementary Fig. S1b). See Sec. 2.6 and Supplementary A.2 for more discussions.

### 2.4 Functional locality-aligned learning yields mechanistically aligned representations

If the functional locality learning paradigm is effective, the learned representations should align with known determinants of enzymatic function. We therefore examined whether the model’s attention patterns reflect catalytic pocket residues, mutational sensitivities, and substrate reaction centers. We first evaluated the EnzymePlex-k_cat_ model on mutant enzyme data from DLKcat [16] (Fig. 4). For two representative enzymes, including polyisoprenyl diphosphate synthase and purine nucleoside phosphorylase, residue-level attention profiles exhibited pronounced peaks concentrated within or proximal to annotated catalytic pocket residues. Wild-type–like mutants predominantly localized to low-attention regions, whereas mutations associated with substantial reductions in k_cat_ were enriched at high-attention positions. These patterns indicate that EnzymePlex captured residue-level functional sensitivity landscapes despite not being trained on mutant data, demonstrating that locality-aligned learning promotes mechanistically meaningful representations rather than superficial correlations.

**Fig. 4:**
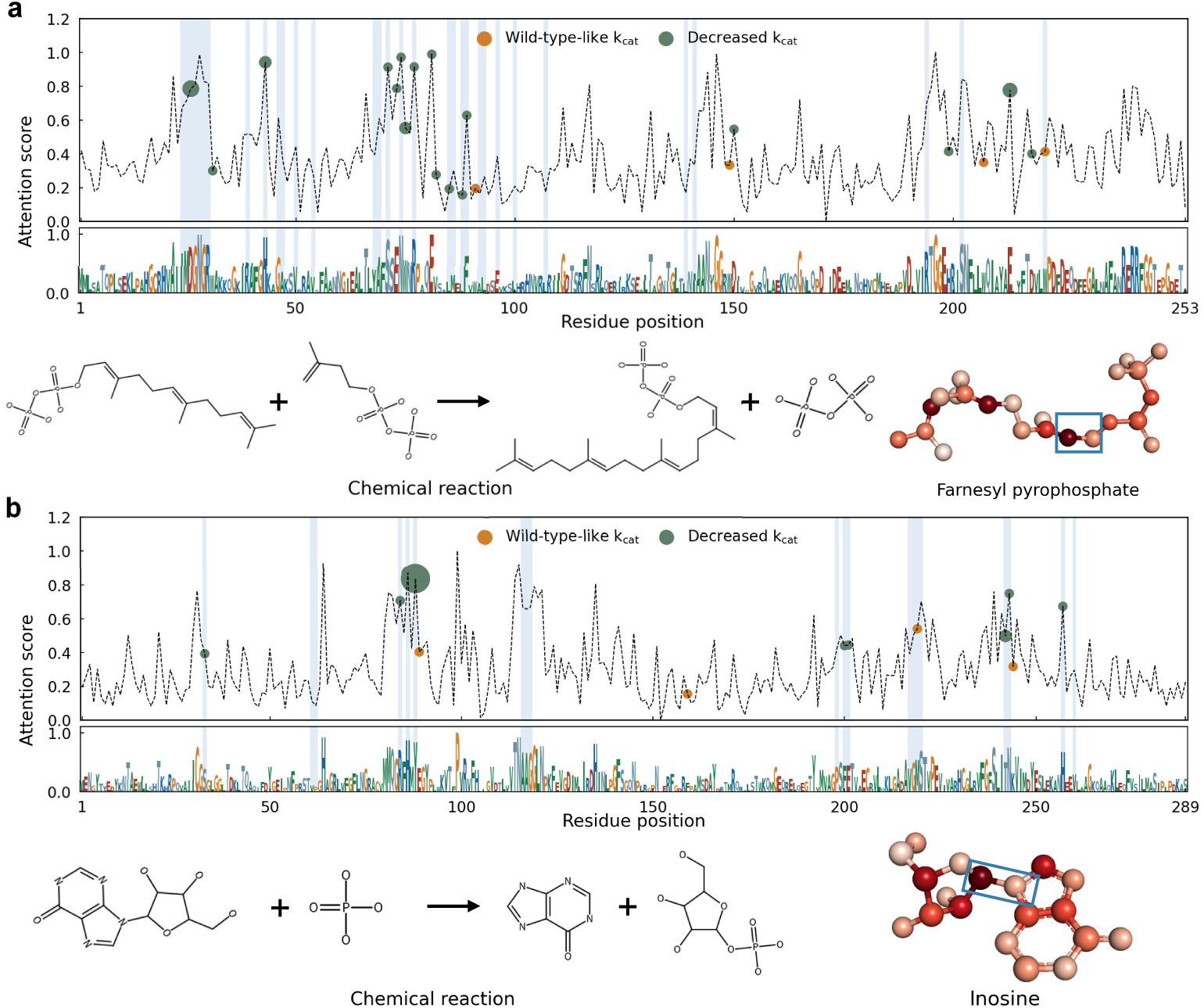
Interpretability analysis of EnzymePlex. For k_cat_ prediction, residue-level attention score profiles of wild-type enzymes are shown. Mutational positions are overlaid on the attention curves, including variants with wild-type-like k_cat_ (**orange** dots) and variants with decreased k_cat_ (**green** dots); dot size indicates the frequency of amino acid substitutions at each position. Sequence logos are generated from residue-wise attention scores of the wild-type enzyme, with letter height proportional to attention magnitude. Catalytic pocket residues are highlighted in **light blue**. Substrate 3D structures are displayed with atom-level attention intensities shown in **red**, and reaction centers outlined by a **blue** bounding box. **(a)** Polyisoprenyl diphosphate synthase (substrate: farnesyl diphosphate; organism: *Escherichia coli* ; EC: 2.5.1.31). **(b)** Purine nucleoside phosphorylase (substrate: inosine; organism: *Homo sapiens*; EC: 2.4.2.1).

Consistent with these residue-level observations, atom-level attention projected onto 3D substrate structures highlighted atoms proximal to annotated reaction centers, which are defined by bond rearrangements, charge shifts, and chirality changes [40]. For farnesyl pyrophosphate (Fig. 4a), elevated attention concentrated on recurring carbon–carbon double-bond motifs, with the highest attention score adjacent to the bond cleavage site; the diphosphate moiety released during catalysis also received increased attention. For inosine (Fig. 4b), atoms surrounding the N-glycosidic bond cleavage site exhibited the strongest attention scores. These patterns demonstrate that EnzymePlex identified chemically meaningful reactive motifs without explicit supervision.

We next quantified modality-level contributions using GradientSHAP [41]. Across all three prediction tasks, EnzymePlex consistently assigned greater importance to catalytic pocket representations than to enzyme sequence representations (Supplementary Fig. S3), underscoring the dominant role of localized pocket structure in determining enzyme kinetics.

### 2.5 Application in synthetic biology and drug discovery

Here, we evaluated the utility of EnzymePlex in two real-world settings: enzyme engineering and small-molecule prioritization. Since published wet-lab measurements often differ substantially from the training distribution (*e.g.*, in temperature, pH, or expression system), our goal is not to predict actual kinetic parameters but to assess whether EnzymePlex could correctly rank enzyme variants by relative catalytic activity and compounds by inhibitory potency, consistent with prior evaluation strategies [8].

#### Enzyme engineering

Tyrosine ammonia lyase (TAL) is a rate-limiting enzyme in the biosynthesis of flavonoids and related industrial compounds [42]. Using wild-type RgTAL as a reference, we evaluated whether EnzymePlex can correctly predict the direction of k_cat_ change for reported mutants. EnzymePlex achieved 66.67% ranking accuracy, whereas CatPred achieved 22.22% (Fig. 5a; Supplementary Fig. S4a). We then assessed single- and triple-point mutants of the enzyme SsCSO from Sphingobium sp. engineered for improved conversion of 4-vinylguaiacol to vanillin [8]. Using the lowest-activity variant (A43N) as a reference, EnzymePlex correctly ranked 9 of 11 variants by k_cat_/K_m_ (81.82%), whereas CatPred achieved only 36.36% accuracy (Fig. 5b; Supplementary Fig. S4b). These results demonstrate the utility of EnzymePlex for prioritizing variants within realistic protein engineering design spaces. **Small-molecule screening.** We next evaluated inhibitor ranking performance.

**Fig. 5:**
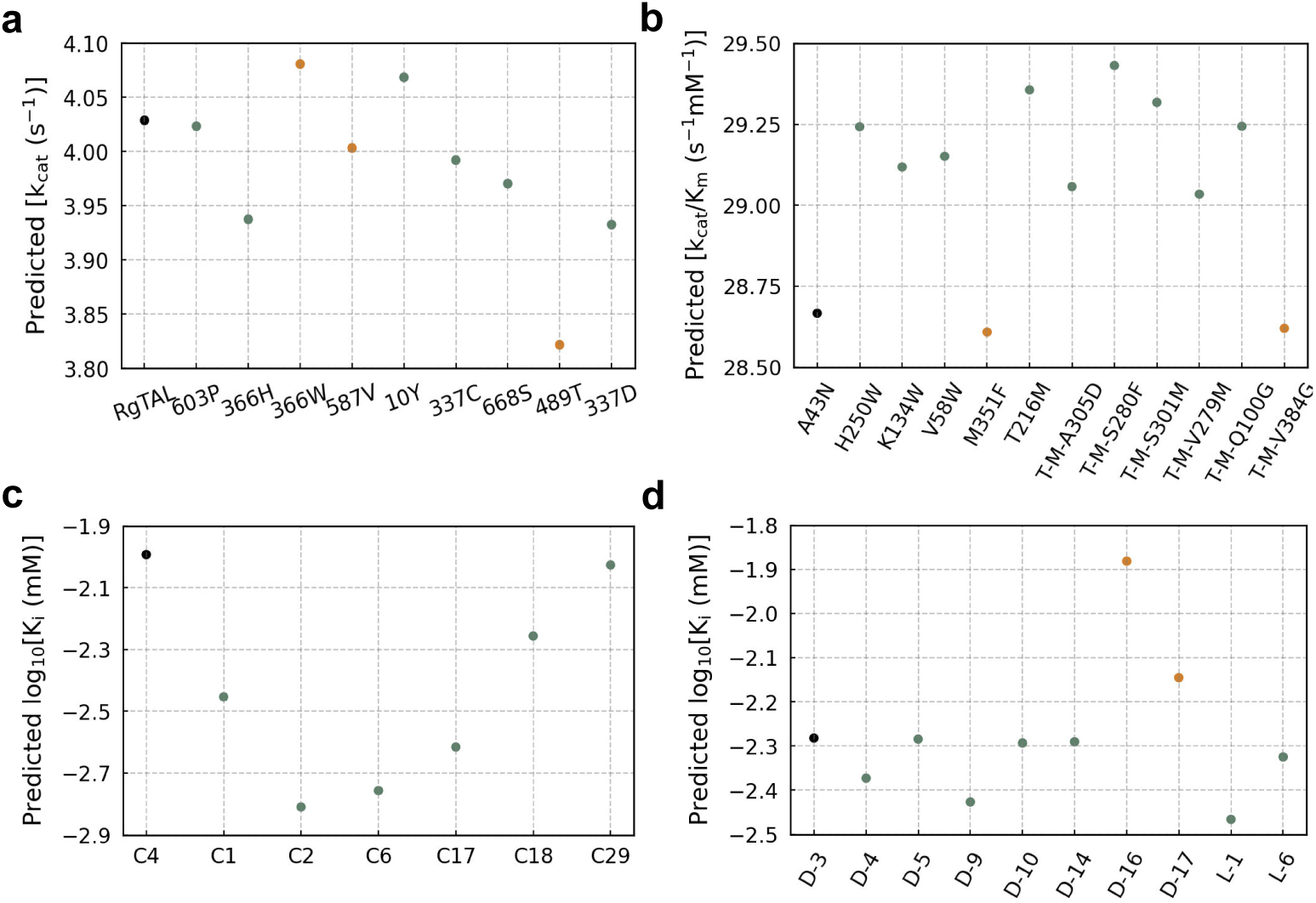
Performance of EnzymePlex on recently reported wet-lab data rel-evant to enzyme engineering and drug discovery. (a) Predicted k_cat_ values for RgTAL mutants studied in flavonoid biosynthesis. (b) Predicted k_cat_/K_m_ values for *Sphingobium* sp. CSO (SsCSO) mutants in carotenoid cleavage oxygenase engineer-ing (“T–M” denotes the M351F–T216M variant). (c) Predicted log_10_[K_i_] values for small-molecule hits targeting the human E3 ligase KLHDC2, a promising platform for PROTAC-based protein degradation. (d) Predicted log_10_[K_i_] values for histone lysine-specific demethylase 1 (LSD1) inhibitors evaluated in virtual screening for non–small cell lung cancer. Black points denote reference samples; **green** points indicate correct predictions, and **orange** points indicate incorrect predictions. A prediction is consid-ered correct if EnzymePlex accurately infers whether catalytic activity or inhibitory potency is higher or lower than the reference.

In a screen targeting the human E3 ubiquitin ligase KLHDC2 [43], which is a KELCH-repeat substrate receptor recently highlighted as a promising PROTAC platform for targeted protein degradation [44]. IC_50_ values were used as qualitative indicators of inhibitory strength, given their correlation with K_i_ [45, 46]. Using the weakest binder (C4) as a reference, EnzymePlex correctly ranked all validated hits, while CatPred correctly ranked four of six compounds (Fig. 5c; Supplementary Fig. S4c). We further analyzed inhibitors of histone lysine-specific demethylase 1 (LSD1) [47, 48], a therapeutic target overexpressed in various solid and hematologic tumors. Among ten active compounds (IC_50_ *<* 100 *µ*M), and using the weakest binder (D-3) as the reference, EnzymePlex achieved 77.78% ranking accuracy and correctly identified L-1 as the most potent inhibitor, whereas CatPred achieved only 22.22% accuracy (Fig. 5d; Supplementary Fig. S4d). Collectively, these results demonstrate that EnzymePlex enables reliable prioritization of enzyme inhibitors under realistic experimental variability, supporting its potential in early-stage drug discovery.

### 2.6 Disentangling the contributions of functional locality, inter-molecule interaction, and substrate geometry

We performed systematic ablations to isolate the contributions of pocket representations, interaction modeling, and substrate geometry. First, removing either pocket structure features or surface features consistently degraded performance (Supplementary Table S1), with larger declines under stringent OOD settings. This indicates that geometric and physicochemical pocket information provide complementary and indispensable features for robust prediction (see Supplementary A.2.1). Second, incorporating global enzyme–molecule interaction improved accuracy relative to non-interaction baselines (rows 2–3 in Fig. 2b and Supplementary Fig. S1b). Importantly, further integrating pocket-level priors into the interaction module yielded additional gains, particularly under severe distribution shift (rows 3–4 in Fig. 2b and Supplementary Fig. S1b), confirming that locality-aligned interaction modeling is critical for generalization (see Supplementary A.2.2). Third, replacing the baseline molecular encoder with a pretrained 3D structure model substantially improved extrapolation to unseen substrates (rows 1–2 in Fig. 2b and Supplementary Fig. S1b). A broader comparison of alternative pretrained molecular encoders, including SMILES-based models [49–51], 2D graph-based models [52–54], and 3D structure–based models [32, 55, 56], is provided in Supplementary A.2.3, which shows that the adopted Uni-Mol2-570M achieves the most satisfactory performance among these alternatives. In addition, substituting ESM-2 with the more recent ESM-3 [57] as the enzyme sequence encoder reduced performance on both test and OOD evaluations (see Supplementary A.2.4), suggesting that advancements in general-purpose pretraining do not necessarily translate to improved task-specific performance. Collectively, these results demonstrate that the robustness of EnzymePlex arises from the synergy of locality-aligned pocket modeling, pocket-guided interaction modeling, and explicit 3D substrate representation.

## 3 Discussion

We have developed EnzymePlex, a functional locality–aligned representation learning framework for predicting enzyme kinetic parameters. Across standard benchmarks, EnzymePlex achieves state-of-the-art performance and exhibits strong generalization to enzymes and substrates that are dissimilar to those encountered during training. Ablation analyses (Sec. 2.6 and Supplementary A.2) demonstrate that these gains arise from explicitly modeling and integrating catalysis-relevant enzyme and substrate structural priors. More broadly, this work exposes a key limitation of prevailing structure-aware learning paradigms: increasing 3D geometric completeness at the global scale does not inherently yield more robust or generalizable function inference. Instead, our findings suggest that the effectiveness of structure-guided learning depends critically on aligning the representational scale with the spatially localized determinants of biological function. This principle may extend beyond enzyme kinetics to other structure-governed biomolecular systems in which function emerges from sparse, localized interactions.

Despite these advances, EnzymePlex currently predicts kinetic parameters solely from enzyme–substrate pairs and does not explicitly incorporate environmental factors such as temperature or pH. Although prior work suggests that including such conditions can improve predictive accuracy [22], progress is constrained by the scarcity of large-scale datasets with high-quality experimental annotations. Expanding enzyme kinetic datasets with richer metadata will therefore be essential for further research. In addition, incorporating experimentally characterized enzyme–substrate poses formed during catalysis represents another promising direction. For non-inhibitory substrates, the catalytically relevant transition-state structures are transient and exist only on femtosecond timescales, making them impossible to isolate or directly characterize experimentally [25]. This motivates exploring approaches such as molecular dynamics simulation methods and generative models designed to produce substrate transition-state structures.

Beyond EnzymePlex’s practical utility in prioritizing high-activity enzymes and high-potency molecular inhibitors for accelerating enzyme engineering and drug discovery, advances in *de novo* protein design [58] further open the possibility of extending EnzymePlex from a predictive tool to a design-oriented framework. When integrated with generative models, such as diffusion-based protein generators [13] and ligand-conditioned sequence generation models [59], EnzymePlex could serve as a property evaluator within closed-loop optimization pipelines, enabling the rational design of enzymes with tailored catalytic behavior.

## 4 Methods

### 4.1 Dataset preparation

For benchmarking and evaluation, we used the CatPred-DB dataset [26], which focuses on kinetic parameters of *wild-type* enzymes and compiles curated measurements of k_cat_, K_m_, and K_i_ from BRENDA [35] (version 2022 2) and SABIO-RK [36] (November 2023). CatPred-DB contains 23,197 k_cat_, 41,174 K_m_, and 11,929 K_i_ entries, each paired with an enzyme amino acid sequence and a substrate SMILES string. All kinetic parameter values were log_10_-transformed. For each prediction task, the data were randomly partitioned into training (80%), validation (10%), and test (10%) sets. For out-of-distribution (OOD) evaluation, we followed the enzyme-level split definitions in the CatPred-DB protocol [26]. Enzyme sequences were clustered using MMseqs2 [60], and test enzymes were selected such that their maximum sequence identity to any enzyme in the training set was below 99%, 80%, 60%, and 40%, respectively. We additionally constructed molecule-level OOD subsets to extend the OOD evaluation beyond enzyme sequence variation and assess the model’s ability to generalize to substrates with low structural similarity. Molecular structures were encoded as Morgan fingerprints [61] generated using RDKit. For multi-component substrates (e.g., “A.B.C”), fingerprints were computed independently for each component and subsequently summed to obtain a composite molecular fingerprint, following [23]. Pairwise molecular similarity was then quantified using the Tanimoto coefficient computed on these aggregated fingerprints. Molecule-level OOD subsets were constructed by selecting only test molecules whose maximum fingerprint similarity to any training molecule was below 99%, 80%, 60%, and 40%, respectively.

We further curated wet-lab–measured data as external test sets to assess the practical utility of EnzymePlex in enzyme engineering and small-molecule inhibitor screening. These include the tyrosine ammonia lyase (TAL) homologue data from Yu et al. [22], the *Sphingobium sp.* CSO mutant data from Wang et al. [8], the KLHDC2 ubiquitin ligase small-molecule screening data (Supplementary Fig. S5) from Zhou et al. [43], and the histone lysine-specific demethylase 1 (LSD1) inhibitor screening data (Supplementary Fig. S6) from Wei et al. [47].

### 4.2 Training and evaluation protocols

EnzymePlex was trained using the negative log-likelihood loss adopted in CatPred [26], which promotes accurate kinetic parameter prediction while regularizing the predicted variance. We optimized all models using AdamW with a learning rate of 1 × 10^−4^ and a batch size of 16. Training was performed on the combined training and validation sets of CatPred-DB [26], and evaluation was carried out on the test set. For fair comparisons, all baseline methods were retrained under identical computational environments with strictly controlled random seeds. Hyperparameter configurations for DLKcat, UniKP, and CatPred followed the CatPred [26] implementation, while those for CataPro [8] were taken directly from its published setup. We adopted a unified evaluation protocol using five independent random seeds, with each seed yielding an ensemble prediction from ten independently trained models. Unless otherwise specified, reported results in Section 2 correspond to the average performance across the five seeds. For the ablation study, each model variant was evaluated using an ensemble of five models trained under the same random seed, and analyses were performed on the CatPred-DB-k_cat_ dataset.

### 4.3 Evaluation metrics

Model performance was assessed using the coefficient of determination (*R*^2^) and the mean absolute error (MAE), two widely used regression metrics that capture complementary aspects of predictive accuracy.

#### Coefficient of determination

(*R*^2^). The *R*^2^ score measures the proportion of variance in the experimental measurements that is explained by the model predictions. Formally, given *y_i_* as the ground-truth kinetic parameter for sample *i* and *y*^*_i_* as the corresponding prediction, *R*^2^ is defined as

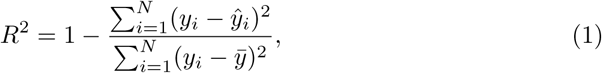

where ̄*y* denotes the mean of the ground-truth values.

#### Mean absolute error (MAE)

MAE quantifies the average magnitude of the prediction error without considering its direction. For *N* samples with predictions *y*^*_i_* and ground-truth values *y_i_*, MAE is given by

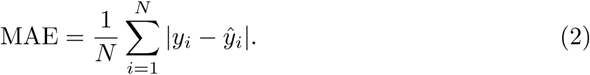

### 4.4 EnzymePlex framework

EnzymePlex takes as input the enzyme’s 3D structure and amino acid sequence, together with the substrate’s 3D conformation, to predict kinetic parameters (Fig. 1a). For each enzyme, we obtained its 3D structure from public databases, using experimental structures when available and AlphaFold models [62] otherwise. Following prior work [40], given an enzyme structure, we first identified the substrate binding pocket using AlphaFill [63], which determines pocket locations through homologous template matching and ligand transfer. If AlphaFill did not produce a confident pocket, we adopted predictions from P2Rank [64]. For each enzyme 3D structure, we extracted both the atomic-level 3D structure of the catalytic pocket and its surface, which together characterize the geometric and physicochemical environment underlying enzyme catalysis. For each substrate, a 3D molecular conformation was generated using RDKit and subsequently refined using the MMFF94 force field. The optimized 3D conformation, together with physicochemical molecular descriptors, was then processed by a pretrained 3D molecular structure encoder to obtain substrate rep-resentations *F*_mol_. The 3D structure, surface topology, and physicochemical properties of the catalytic pocket were encoded with an SE(3)-invariant graph neural network, producing pocket-structure features 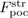 and pocket-surface features 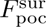. In parallel, the enzyme sequence was encoded using the pretrained protein language model, yielding sequence representations *F*_seq_.

To capture enzyme-substrate interaction patterns, EnzymePlex employs a pocket-guided multi-modal interaction module that models the interplay among the substrate molecule, the catalytic pocket, and the enzyme sequence. We denote the resulting catalysis-aware representations as 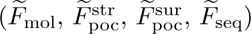. These representations are subsequently passed through attention pooling layers, AttPool. For an input feature matrix *F* ∈ ℝ*^L^*^×*C*^, attention pooling computes a scalar score for each position,

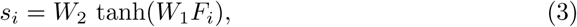

and computes a weighted aggregation of the input features:

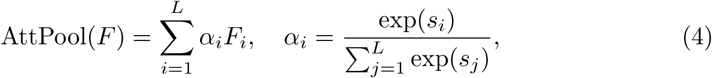

where *W*_1_ ∈ ℝ*^H^*^×*C*^ and *W*_2_ ∈ ℝ^1×*H*^ are learnable parameters and tanh(·) introduces nonlinearity. This formulation enables EnzymePlex to adaptively highlight structurally or functionally informative representations during feature aggregation. Finally, the pooled representations from all modalities are concatenated and passed through a four-layer MLP to predict the kinetic parameter *k*:

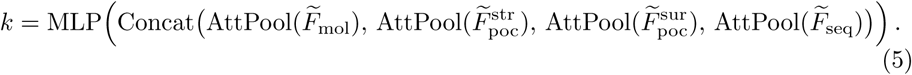

#### 4.4.1 Catalytic pocket structure and surface extraction

Enzyme kinetics are strongly shaped by enzyme–substrate interactions within the catalytic pocket, where substrates bind and undergo catalysis. To model these functionally critical regions, we explicitly extracted the 3D structure of the catalytic pocket. Inspired by Liu et al. [40], we applied AlphaFill [63] to transfer ligand information from homologous protein–ligand complexes in the PDB-REDO database [65] to each query structure through template matching and structural alignment, thereby identifying high-confidence catalytic pockets. If AlphaFill did not produce a valid pocket, we instead used P2Rank [64], a machine-learning-based pocket prediction method, and retained the two pockets with the highest confidence scores.

The catalytic pocket surface constitutes the immediate physical interface at which the substrate contacts the enzyme. To characterize this interface, we employed MaSIF [66] to compute surface interaction fingerprints for each pocket. These fingerprints encode geometric descriptors such as shape index and distance-dependent curvature, together with chemical descriptors including hydrophobicity, hydrogen-bonding potentials, and Poisson–Boltzmann electrostatics, providing a detailed representation of the microenvironment relevant to substrate recognition and catalytic activity.

#### 4.4.2 SE(3)-invariant pocket encoder

We modeled catalytic pockets using an SE(3)-invariant graph neural network that performs rotation- and translation-invariant message passing over pocket structures and surfaces. We first constructed a residue-level pocket structure graph *G*^str^ = (*V* ^str^*, E*^str^), with node positions defined by residue C*_α_* coordinates. Each node was annotated with SE(3)-invariant features, including amino-acid identity, local self-distance descriptors, backbone dihedral angles, and ESM-2 [19] embeddings. Residue–residue edges were added when the minimum atom–atom distance between residue pairs was below a 15 Ä cutoff. Edge attributes comprised a backbone-connectivity indicator, several distance-based geometric descriptors, and Gaussian radial-basis expansions of the C*_α_*–C*_α_*distance. We then constructed a pocket surface graph *G*^sur^ = (*V* ^sur^*, E*^sur^) from MaSIF-derived surface meshes [66]. Each surface vertex served as a node and was annotated with hydrophobicity, hydrogen-bonding potentials, Poisson–Boltzmann electrostatics, shape index, and curvature. Edges were defined by mesh connectivity between adjacent vertices, and edge attributes consisted of Gaussian radial-basis encodings of pairwise surface-vertex distances. To integrate these two modalities, we additionally constructed a pocket structure–surface interaction graph *G*^str^ ^sur^ = (*V* ^str^ ^sur^*, E*^str^ ^sur^). Each receptor residue (C*_α_* position) was connected to nearby surface vertices within a 15 Ä radius, with at most 30 surface neighbors per residue. Node features were inherited directly from the structure and surface graphs, and edge attributes were defined as Gaussian radial-basis encodings of distances between surface vertices and residue C*_α_* positions.

All three pocket graphs were processed using stacked SE(3)-invariant graph convolution layers equipped with residual connections. Let *i* be a node and *j* ∈ N (*i*) a neighboring node, with *h_i_* and *e_ij_* denoting their node and edge features. Each layer updates node representations via

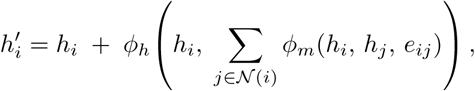

where *ϕ_m_* and *ϕ_h_* are learnable MLPs. The residual pathway preserves information across layers and contributes to more stable optimization during message passing.

We applied four such layers to the pocket structure graph, followed by two layers to the pocket structure–surface interaction graph, and four layers to the pocket surface graph. This hierarchical processing flow first aggregates residue-level geometric representations, then propagates structural context to surface vertices, and finally refines the surface representations with localized surface–surface interactions. The hidden dimensionality was set to 128 for the k_cat_ and K_m_ tasks, and reduced to 64 for the K_i_ task to mitigate overfitting given the smaller size of the K_i_ dataset. The resulting node embeddings from the structure and surface graphs form the pocket-structure features *F* ^str^ and pocket-surface features *F* ^sur^, which jointly capture the geometric and physicochemical characteristics of each catalytic pocket.

#### 4.4.3 3D molecular structure encoder

To evaluate the impact of molecular representation choice on model performance, we compared a diverse set of pretrained small-molecule encoders spanning multiple input modalities. These included SMILES-based models (SMILES-TRANS [20] and SMILES-BERT [50]), 2D graph-based models (MolCLR [52], GROVER [53] and GraphMVP [54]), and 3D structure–based models (GEM [55], Uni-Mol [56] and Uni-Mol2 [32]).

Across these comparisons, the Uni-Mol family consistently demonstrates the strongest molecular-level OOD generalization while also exhibiting solid enzyme-level generalization performance (Supplementary Table S2). Based on this observation, we adopted Uni-Mol2-570M as the default 3D molecular structure encoder in EnzymePlex and used its atom-level outputs as the molecular features *F*_mol_.

#### 4.4.4 Enzyme sequence encoder

Pretrained self-supervised protein language models have been widely adopted for enzyme function prediction [12, 40] and kinetic parameter modeling [8, 26]. We adopted ESM2 [19] as our default sequence encoder to ensure fair comparison with Cat-Pred [26]. ESM-2 is trained with a masked language modeling objective that predicts the identities of randomly masked amino acids within a protein sequence. We employed the 650M-parameter variant of ESM-2, which produces 1280-dimensional residue-level embeddings. Residue-level representations *F*_seq_ are extracted from the final hidden layer and subsequently passed to the downstream modules of EnzymePlex. In addition, we also employed the open-source ESM-3-open [57] model (1.4B parameters), which produces 1536-dimensional residue-level representations.

#### 4.4.5 Pocket-guided multi-modal interaction module

We introduced a pocket-guided multi-modal interaction module that integrates substrate, pocket, and sequence representations through an asymmetric cross-attention mechanism [67]. The asymmetry arises from the distinct roles assigned to each modality in the cross-attention mechanism. Specifically, catalytic pocket information is injected into both molecular and sequence representations; substrate molecular information is injected into both sequence and pocket representations; whereas enzyme sequence information is injected only into molecular representations. The pocket-guided multi-modal interaction module comprises three components: molecule self-adaptation, global enzyme-molecule interaction, and pocket-prior guided local interaction.

##### Molecule self-adaptation

We first adapted the molecular features *F*_mol_ using a multi-head self-attention [67] layer:

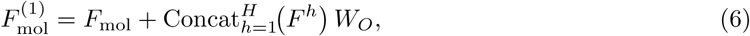

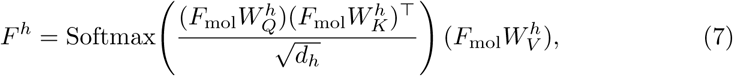

where 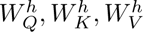 are the projection matrices for head *h*, *d_h_* is the dimensionality of each head, and *W_O_* is the output projection. This self-attention mechanism captures intramolecular dependencies and produces task-adaptive substrate representations.

##### Global enzyme–molecule interaction

We modeled global enzyme-molecule interactions by applying bi-directional multi-head cross-attention between the full enzyme sequence and the substrate molecule. The cross-attention operator [67] is defined as

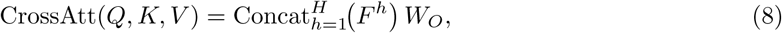

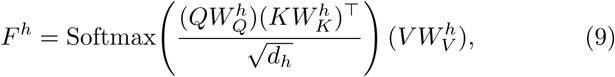

where *H* is the number of attention heads, *d_h_*is the dimensionality of each head, and 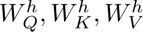 are learnable projection matrices.

Molecule-to-sequence attention is formulated as

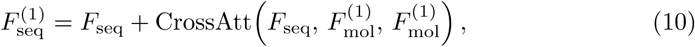

and sequence-to-molecule attention is given by

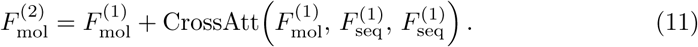

Together, this reciprocal information exchange allows enzyme sequence representations to incorporate molecule-induced biochemical cues, while molecular representations capture enzyme-specific functional context.

##### Pocket-prior guided local interaction

We further refined sequence and molecular representations using localized pocket surface features, which aggregate geometric and physicochemical properties across the catalytic pocket. At the same time, the pocket structural representation 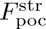 is passed through the module unchanged via an identity mapping, producing 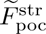.

Pocket-to-sequence information injection is implemented as:

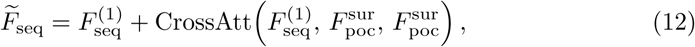

allowing enzyme sequence representations to perceive localized structural features of the catalytic pocket.

Pocket-molecule interactions are then modeled via:

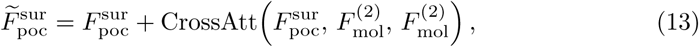

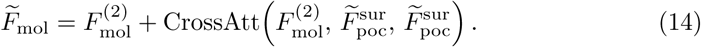

Through these pocket-guided interactions, enzyme sequence and molecular representations are explicitly informed by localized geometric and physicochemical patterns of the catalytic pocket, while pocket features concurrently incorporate substrate-aware information.

### 4.5 Biochemical pattern analyses

To assess whether EnzymePlex captures established biochemical patterns, we conducted three analyses: enzyme metabolism patterns, enzyme promiscuity, and substrate-associated enzyme diversity. For metabolism-related patterns, enzymes were assigned to either primary central and energy metabolism or intermediary and secondary metabolism based on their metabolic context. Predicted kinetic parameters (k_cat_, K_m_, K_i_) were independently compared between these groups to evaluate whether EnzymePlex reflects known metabolic trends. For enzyme promiscuity, substrates of promiscuous enzymes (*i.e.*, enzymes acting on multiple substrates) were partitioned into preferred and alternative categories based on experimentally measured kinetics.

For k_cat_, the substrate with the highest measured value was designated as preferred, with all others classified as alternative; for K_m_, the substrate with the lowest measured value was designated as preferred, with the remaining substrates classified as alternative. Predicted k_cat_ and K_m_ values were analyzed separately to assess whether EnzymePlex captures substrate preference. Entries with experimentally measured log_10_[k_cat_] less than 2 (s^−1^) and log_10_[K_m_] more than 2 (mM) were excluded from this analysis. For substrate-associated enzyme diversity, enzymes linked to the same substrate were grouped into native and underground categories based on experimental K_i_ measurements. The enzyme with the lowest measured K_i_ was designated as native, and the remaining enzymes were classified as underground. Entries with experimentally measured log_10_[K_i_] more than 2 (mM) were excluded. All statistical comparisons were performed using two-sided Wilcoxon rank-sum tests.

### 4.6 Interpretation analyses

#### Fine-grained residue and atom attribution analysis

To understand how EnzymePlex prioritizes residues and substrate atoms during k_cat_ prediction, we analyzed the attention scores *α_i_*(Eq. 4) produced by the attention pooling layer when applied to the enzyme and molecule representations. These scores indicate the importance of each residue or atom.

For visualization, the raw scores were further min–max normalized on a per-sample basis to emphasize relative importance along enzyme sequences or substrate atoms. The resulting normalized profiles were then used to plot residue-level attention score curves and to map atom-level attention values onto the 3D substrate structures. Together, these visualizations provide fine-grained insights into the catalytic determinants captured by EnzymePlex.

#### Multi-modal feature attribution using GradientSHAP

We used Gradi-entSHAP [41, 68] to quantify the contribution of each modality. For every test sample, we extracted intermediate enzyme sequence, enzyme pocket, and molecular features using a feature-only forward pass, and applied GradientSHAP with Gaussian noise baselines to obtain attribution maps. The absolute attributions were averaged within each modality and then aggregated across all test samples. The resulting modal contributions were normalized, reweighted by *d^α^* (*α* = −0.5) to adjust for different feature dimensionalities, and renormalized. Reported values were averaged across ten ensemble models.

## Data and model availability

Training and testing data for k_cat_, K_m_, and K_i_ were obtained from the open-source CatPred-DB benchmark [https://doi.org/10.5281/zenodo.14775076]. The molecule-level out-of-distribution (OOD) subsets constructed in this study are publicly available at [https://github.com/zhanghao5201/EnzymePlex]. The mutant-enzyme data used for the interpretation analyses were collected from [https://github.com/SysBioChalmers/DLKcat]. The tyrosine ammonia lyase homologue data was obtained from [https://doi.org/10.1038/s41467-023-44113-1]. The *Sphingobium sp.* CSO mutant data was obtained from [https://doi.org/10.1038/s41467-025-58038-4].

The human KLHDC2 ubiquitin ligase small molecule screening data was obtained from [https://doi.org/10.1038/s41467-024-52061-7]. The histone lysine-specific demethylase 1 inhibitor screening data was obtained from [https://doi.org/10.1038/s41401-024-01439-w]. PDB-REDO is available at [https://pdb-redo.eu/]. The trained model checkpoints are available at GitHub [https://github.com/zhanghao5201/EnzymePlex].

## Code availability

The source code for EnzymePlex is available at Github [https://github.com/zhanghao5201/EnzymePlex].

## Acknowledgements

This work is supported by the Fundamental and Interdisciplinary Disciplines Break-through Plan of the Ministry of Education of China under Grant JYB2025XDXM504. He Zhang is supported by the Zhongguancun Academy under Grant No.C20250512.

## Author contributions

N.Z. and H.Z. conceived and supervised the research. Ha.Z., H.Z., M.K., K.Z., T.Y., and N.Z. contributed to the methodology design and result analyses. Ha.Z. curated the data. Ha.Z., H.Z., and K.Z. contributed to data curation, algorithm implementation, and model training. H.Z., Ha.Z., N.Z., and M.K. wrote the manuscript with the input from other authors. All authors were involved in the discussion and proofreading.

## Inclusion & ethics

All contributors who fulfill the authorship criteria are listed as co-authors in this paper.

## Appendix A Supplementary Material

**Fig. S1:**
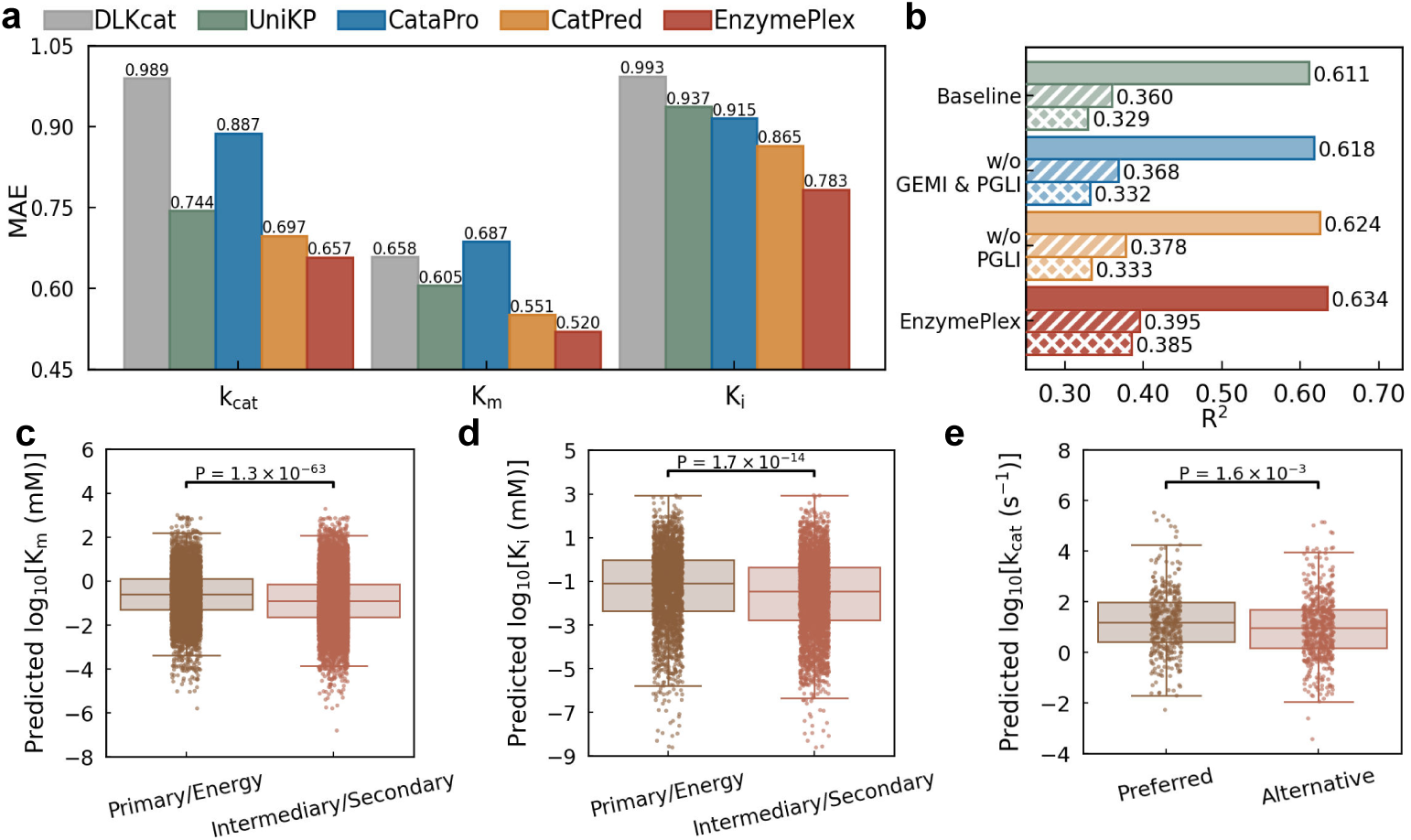
Performance evaluation of EnzymePlex, as an extension to the results narrated in Sec. 2.2. **(a)** Benchmark comparison of EnzymePlex with other methods on the CatPred-DB test set. Mean absolute error (MAE) values are reported. **(b)** *R*^2^ results of EnzymePlex variants for k_cat_ prediction on the test set (solid bars) and enzyme-level OOD subsets. **(c)** Comparison of EnzymePlex performance on K_m_ for enzymes involved in primary central and energy metabolism versus intermediary and secondary metabolism. **(d)** Comparison of EnzymePlex performance on K_i_ for enzymes involved in primary central and energy metabolism versus intermediary and secondary metabolism. **(e)** Enzyme promiscuity analysis for k_cat_. For enzymes acting on multiple substrates, substrates were grouped into preferred and alternative cate-gories based on experimentally measured kinetics. Two-sided Wilcoxon rank-sum tests were used to compute *P* values. The central line denotes the median, the box shows the interquartile range, and whiskers extend to 1.5 times the interquartile range.

### A.1 Additional out-of-distribution generalization results

As shown in Sec. 2.3, EnzymePlex exhibits substantial improvement on out-of-distribution (OOD) evaluation in terms of *R*^2^. Here we demonstrate that MAE reflects performance advantages similar to those observed with *R*^2^ across both enzyme-level and molecule-level OOD subsets, as shown in Supplementary Fig. S2. An exception occurred for K_m_ at the most challenging 40% similarity split, where EnzymePlex (MAE = 0.690) underperformed UniKP (MAE = 0.680) and showed improvement over CatPred (MAE = 0.709). This discrepancy likely reflects differences in metric sensitivity under extreme OOD conditions. MAE measures absolute prediction error and can therefore be disproportionately influenced by a small number of large deviations, particularly when kinetic parameters span multiple orders of magnitude. By contrast, *R*^2^ evaluates the proportion of variance explained and better captures whether global trends in the data are preserved, under which EnzymePlex achieved improved performance.

**Fig. S2:**
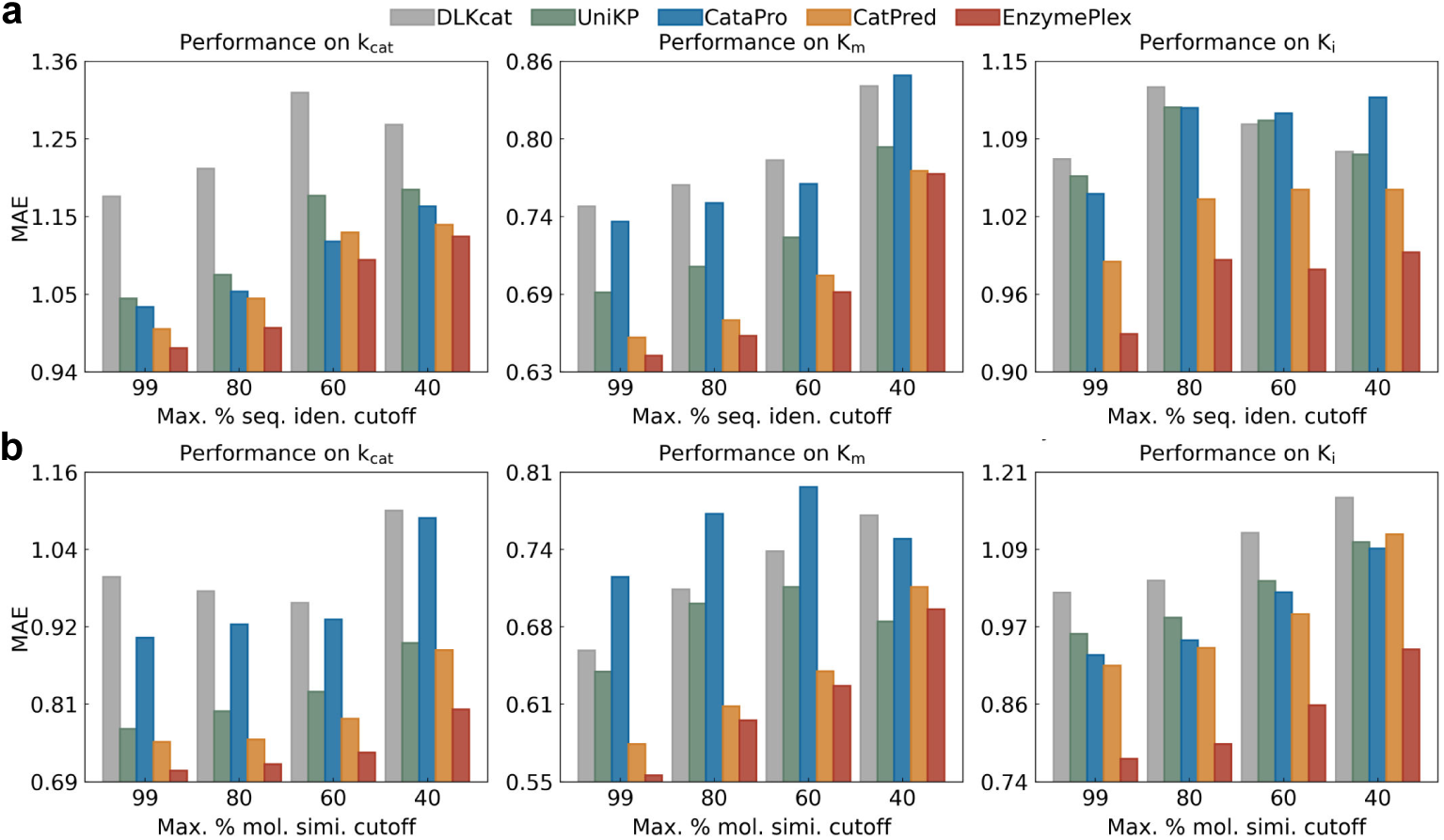
Out-of-distribution performance of EnzymePlex compared with other methods, as an extension to the results narrated in Sec. 2.3. MAE values are reported. **(a)** Enzyme-level OOD performance under sequence identity cut-offs. **(b)** Molecule-level OOD performance under molecular similarity cutoffs.

**Table S1:**
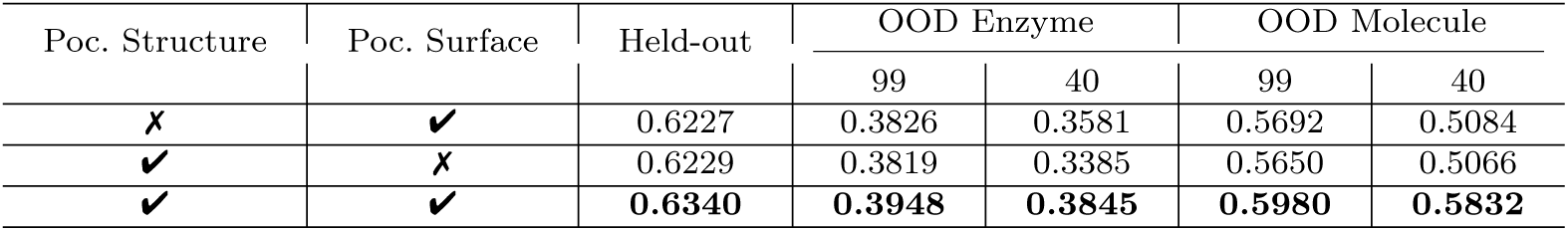
Performance comparison of enzyme pocket encoders on the CatPred-DB-k_cat_ dataset for OOD generalization. Uni-Mol2-570M [32] is used as the molecule encoder, and ESM2-650M [19] is used as the enzyme sequence encoder. *R*^2^ is reported as the evaluation metric. The OOD enzyme subsets include only enzymes whose maximum sequence identity to any training sequence is below the indicated thresholds (99% and 40%). The OOD molecule subsets include only molecules whose maximum molecular fingerprint similarity to any training molecule is below the corresponding thresholds (99% and 40%).

**Fig. S3:**
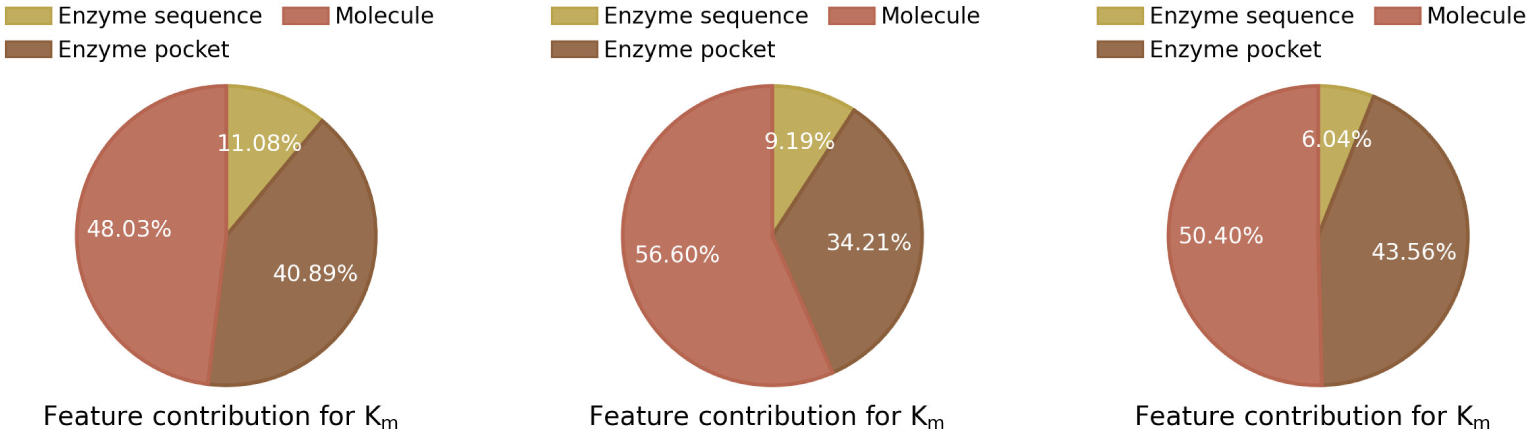
Feature contribution analysis across modalities for k_cat_, K_m_, and K_i_ prediction on the test set, as an extension to the results narrated in Sec. 2.6. Feature attributions are computed using GradientSHAP for sequence, pocket, and molecular features. For each modality, absolute attribution scores are averaged across all input samples, and all resulting contributions from enzyme sequence, enzyme pocket, and substrate features are normalized to yield the relative proportions displayed in the pie charts.

**Table S2:**
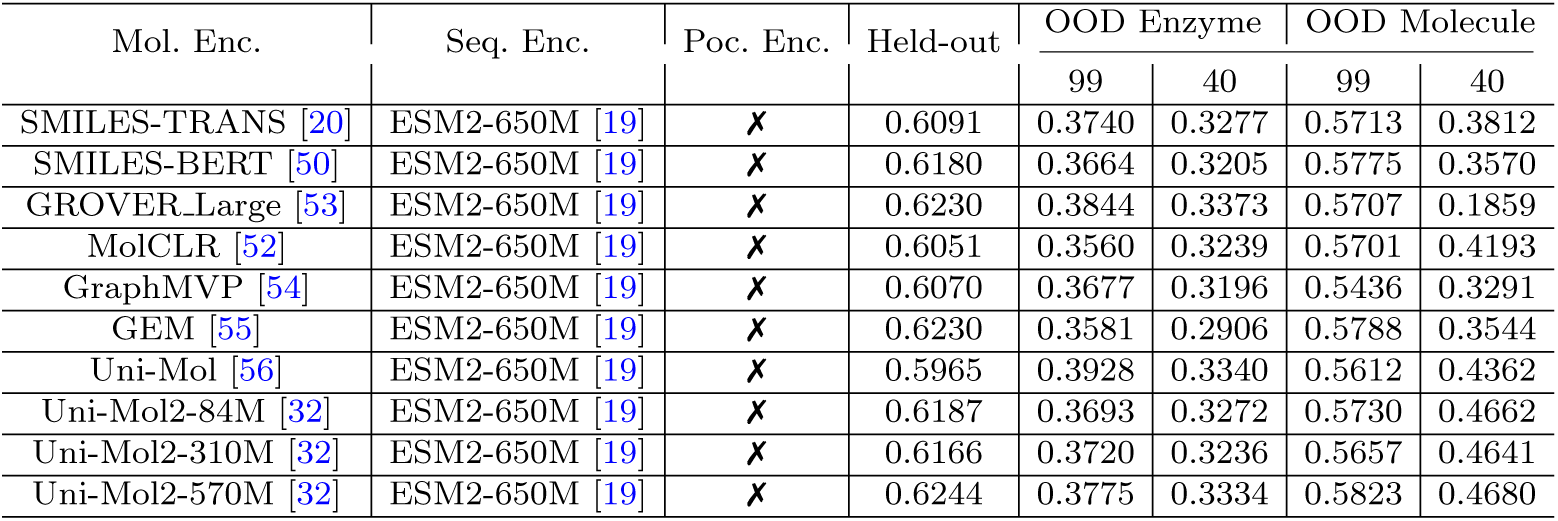
Performance comparison of pretrained molecule encoders on the CatPred-DB-k_cat_ dataset for OOD generalization. *R*^2^ is reported as the evaluation metric.

### A.2 Disentangling the contributions of functional locality, inter-molecule interaction, and substrate geometry

#### A.2.1 Contribution of pocket structure and surface representations

To examine the respective contributions of pocket structural and pocket surface representations, we selectively removed each component from both the pocket encoder and the pocket-guided multi-modal interaction module. As shown in Supplementary Table S1, removing pocket surface modeling while retaining pocket structure modeling resulted in a noticeable performance degradation, with the *R*^2^ on the test set decreasing to 0.6229. Performance also declined on the molecule-level OOD test sets by 5.52% and 13.13% at the 99% and 40% similarity cutoffs, respectively, and on the enzyme-level OOD test sets by 3.27% and 11.96% at the corresponding cutoffs. Con-versely, removing pocket structure modeling while retaining surface modeling yielded an *R*^2^ of 0.6227 on the test set, accompanied by performance reductions of 4.82% and 12.83% on the molecule-level OOD test sets at the 99% and 40% cutoffs, respec-tively, and decreases of 3.09% and 6.87% on the enzyme-level OOD test sets. Taken together, these results indicate that both pocket structure and surface representations contribute meaningfully to enzyme kinetic parameter prediction, and that the struc-tural and physicochemical information captured by these components is essential for achieving accurate and generalizable performance.

#### A.2.2 Necessity of pocket-guided multi-modal interaction

We next evaluated the respective contributions of global enzyme-molecule interaction (GEMI) and pocket-prior guided local interaction (PGLI) within the pocket-guided multi-modal interaction module. As shown in Fig. 2b, incorporating GEMI led to clear performance improvements over the model variant without any enzyme-molecule interaction, increasing *R*^2^ by 1.57% and 3.54% on the molecule-level OOD test sets at the 99% and 40% similarity cutoffs, respectively, and by 2.72% on the enzyme-level OOD test set at the 99% cutoff (Supplementary Fig. S1b). These results demonstrate the effectiveness of modeling global enzyme-molecule interactions through enzyme sequence and molecular features. When pocket priors were further incorporated into the interaction module (PGLI), EnzymePlex achieved additional and substantially larger performance gains. Specifically, *R*^2^ increased by 2.75% and 24.57% on the molecule-level OOD test sets at the 99% and 40% similarity cutoffs, respectively (Fig. 2b), and by 4.50% and 15.62% on the enzyme-level OOD test sets at the corresponding cutoffs (Supplementary Fig. S1b). These gains indicate that integrating pocket-level structural priors enables the model to capture catalytically relevant local interaction patterns, thereby improving both predictive accuracy and generalization of enzyme kinetic parameters.

#### A.2.3 Effect of the pretrained 3D molecular structure encoder

We assessed the contribution of the pretrained 3D molecular structure encoder by replacing the baseline 2D graph-based molecular encoder MPNN [38] with Uni-Mol2-570M [32]. This replacement led to clear improvements in molecular generalization. Specifically, the resulting variant achieved an *R*^2^ of 0.618 on the k_cat_ test set and exhibited performance gains of 3.80% and 13.85% on the molecule-level OOD test sets at the 99% and 40% similarity cutoffs, respectively (Fig. 2b). The corresponding improvements under enzyme-level OOD settings were comparatively modest (Supplementary Fig. S1b), indicating that incorporating explicit substrate 3D structure modeling primarily benefits generalization to previously unseen molecules. A broader comparison across diverse pretrained molecular representation models, including SMILES-based models [49–51], 2D graph-based models [52–54], and 3D structure–based models [32, 55, 56], further showed that the Uni-Mol2 family consistently achieved the strongest molecular OOD generalization while maintaining competitive performance on both the test set and enzyme-level OOD evaluations (Supplementary Table S2). These results highlight the substantial advantage gained from explicitly modeling 3D molecular geometry in enhancing generalization to unseen molecules.

#### A.2.4 Comparison of protein language model variants

To remain consistent with CatPred [26], we used ESM-2 [19] as the default protein sequence encoder. We further conducted an exploratory comparison by replacing ESM-2 with the more recent ESM-3 model [57]. As shown in Supplementary Table S3, this substitution led to reduced performance on both the test set and the OOD evaluations. A similar observation was reported in a recent study [14], where ESM-3 did not out-perform ESM-2 on downstream tasks requiring fine-grained sequence discrimination. This may reflect differences in pretraining objectives: ESM-3 is optimized for multi-modal generative modeling across sequence, structure, and function, whereas ESM-2’s masked-token pretraining is more directly aligned with capturing residue-level variation relevant to enzyme kinetic prediction. These findings suggest that applying large protein language models to enzyme kinetic parameter prediction remains an open chal-lenge, and that further advances in sequence representation learning are needed for this task.

**Table S3:** Performance comparison of pretrained enzyme sequence encoders on the CatPred-DB-k_cat_ dataset for OOD generalization. *R*^2^ is reported as the evaluation metric.

| Mol. Enc. | Seq. Enc. | Poc. Enc. | Held-out | OOD Enzyme |  | OOD Molecule |  |
| --- | --- | --- | --- | --- | --- | --- | --- |
|  |  |  |  | 99 | 40 | 99 | 40 |
| Uni-Mol2-570M [32] | ESM2-650M [19] | ✓ | **0.6340** | **0.3948** | **0.3845** | **0.5980** | **0.5832** |
| Uni-Mol2-570M [32] | ESM3-open [57] | ✓ | 0.6168 | 0.3548 | 0.3033 | 0.5835 | 0.5063 |

**Fig. S4:**
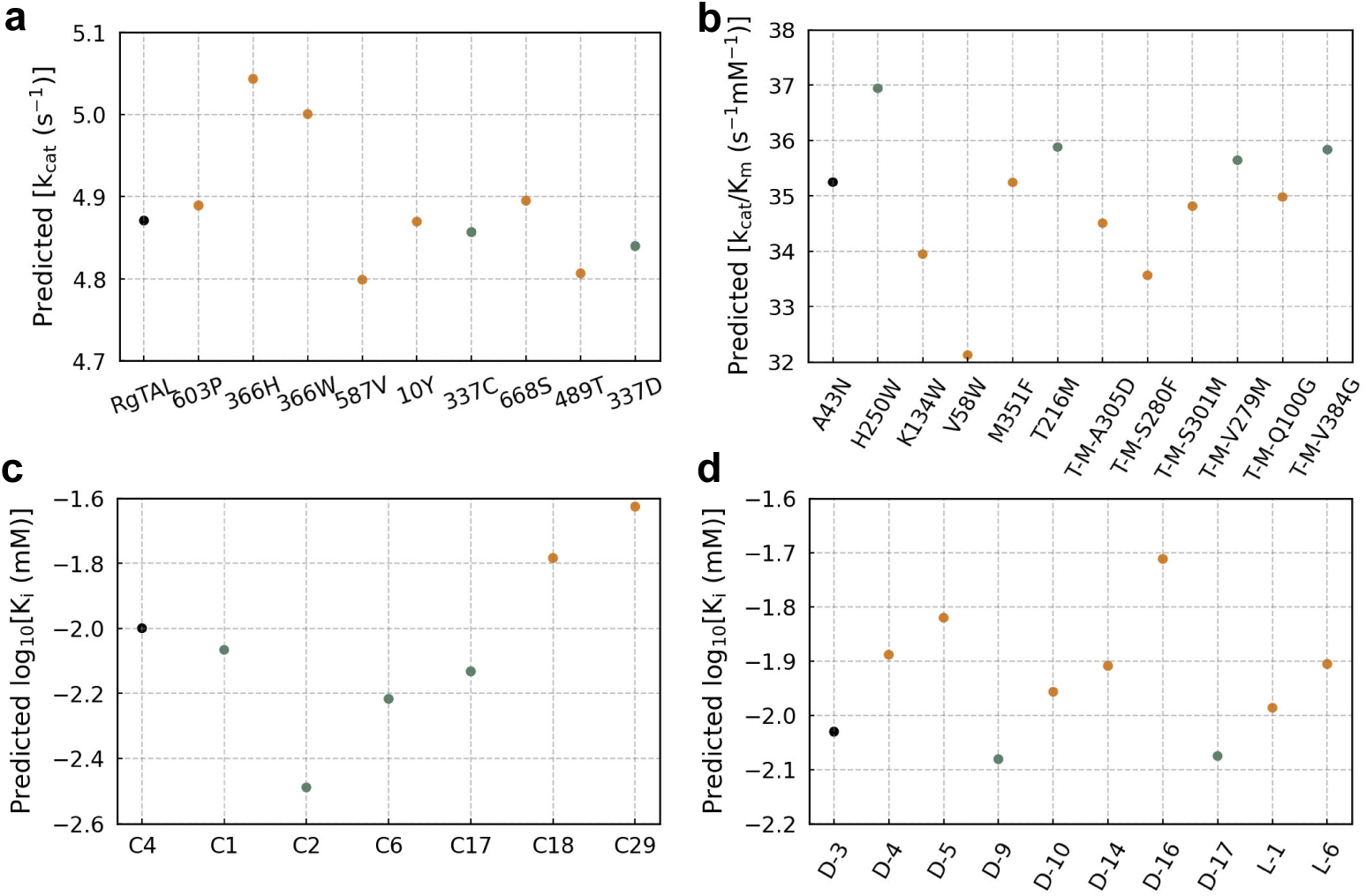
Performance of CatPred on recently reported wet-lab data rele-vant to enzyme engineering and drug discovery. **(a)** Predicted k_cat_ values for RgTAL mutants used in flavonoid biosynthesis research. **(b)** Predicted k_cat_/K_m_ values for *Sphingobium* sp. CSO (SsCSO) mutants involved in carotenoid cleavage oxygenase mining and engineering (“T–M” denotes the M351F–T216M variant). **(c)** Predicted log_10_[K_i_] values for small-molecule hits targeting the human E3 ligase KLHDC2, apromising platform for PROTAC-based targeted protein degradation. **(d)** Predicted log_10_[K_i_] values for histone lysine-specific demethylase 1 (LSD1) inhibitors evaluated in virtual-screening studies for potential therapeutic applications in non–small cell lung cancer. Black points denote reference samples. **Green** points indicate correct predic-tions, whereas **orange** points indicate incorrect predictions. For **(a–d)**, a prediction is considered correct if EnzymePlex correctly infers whether the activity or inhibitory strength of a given sequence or molecule is higher or lower relative to the reference sample.

**Fig. S5:**
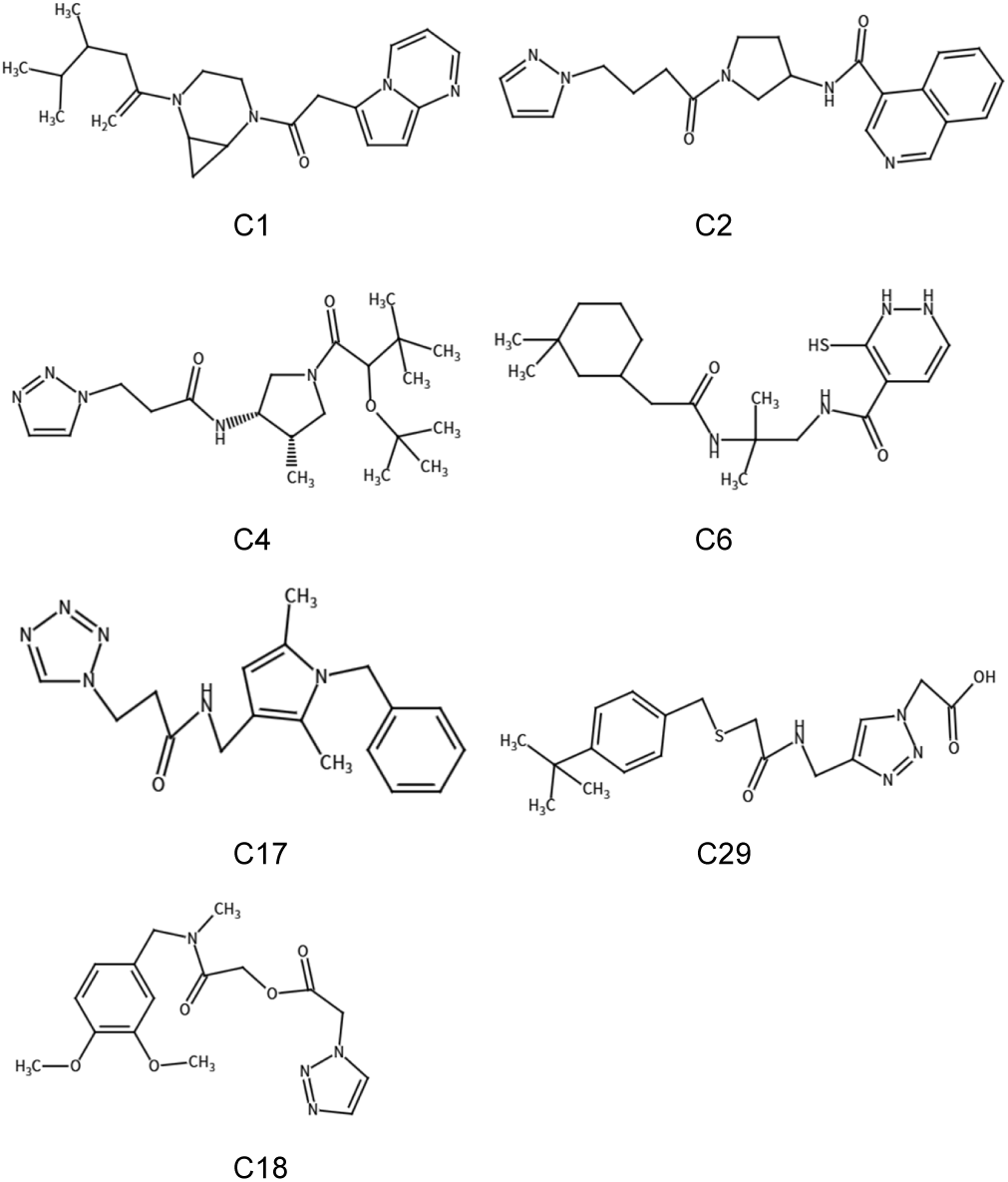
Small molecules from the KLHDC2 ubiquitin ligase screening dataset.

**Fig. S6:**
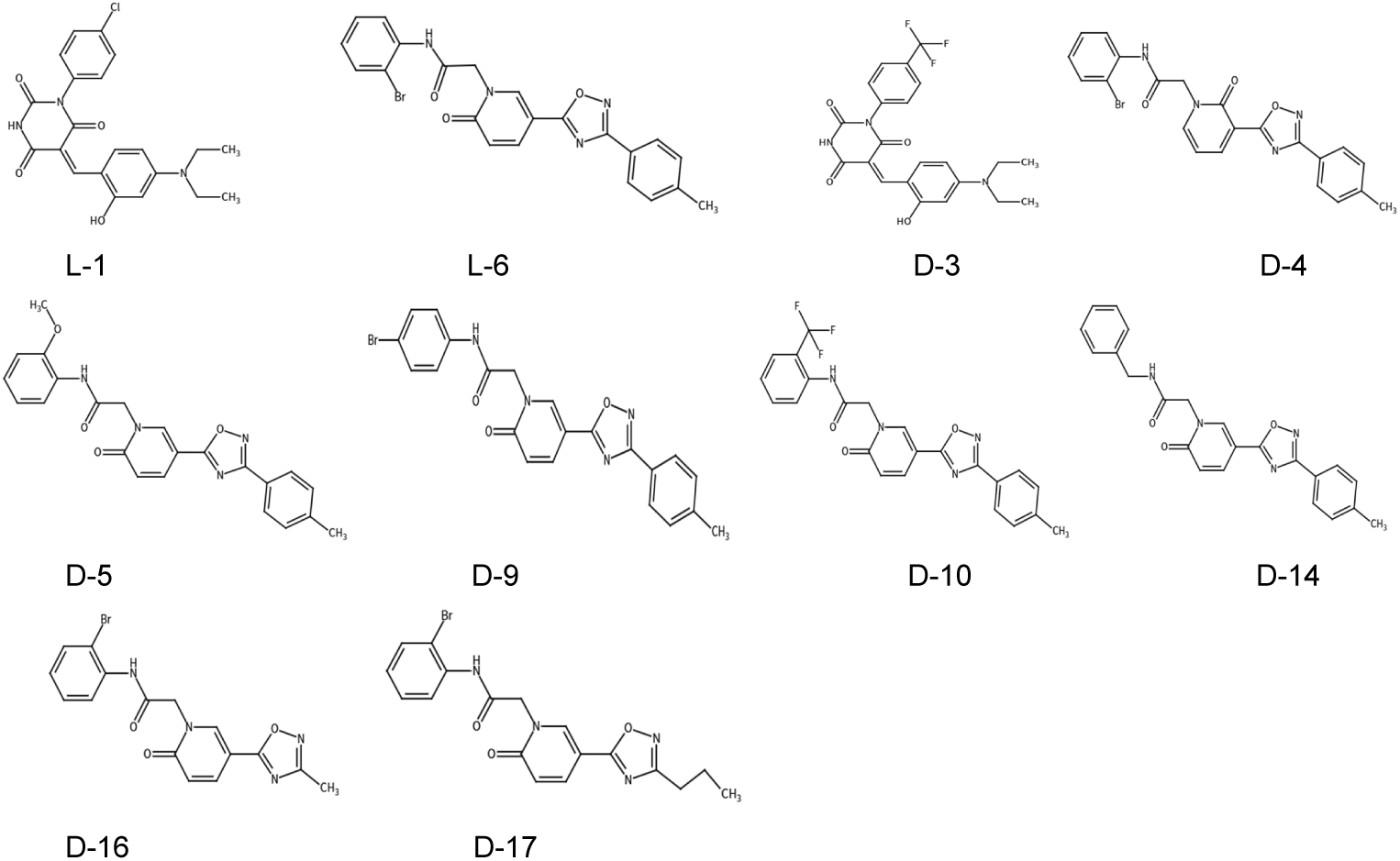
Small molecules from the histone lysine-specific demethylase 1 (LSD1) inhibitor screening dataset.

## References

[1] Benítez-Mateos, A.I., Roura Padrosa, D., Paradisi, F.: Multistep enzyme cascades as a route towards green and sustainable pharmaceutical syntheses. Nature Chemistry 14(5), 489–499 (2022)

[2] Landwehr, G.M., Bogart, J.W., Magalhaes, C., Hammarlund, E.G., Karim, A.S., Jewett, M.C.: Accelerated enzyme engineering by machine-learning guided cell-free expression. Nature Communications 16(1), 865 (2025)

[3] Oliveira, J.T., Costa, F.M., Silva, T.G., Simöes, G.D., Santos Pereira, E., Costa, P.Q., Andreazza, R., Schenkel, P.C., Pieniz, S.: Green tea and kombucha characterization: Phenolic composition, antioxidant capacity and enzymatic inhibition potential. Food Chemistry 408, 135206 (2023)

[4] Buller, R., Lutz, S., Kazlauskas, R., Snajdrova, R., Moore, J., Bornscheuer, U.: From nature to industry: Harnessing enzymes for biocatalysis. Science 382(6673), 8615 (2023)

[5] Bond-Watts, B.B., Bellerose, R.J., Chang, M.C.: Enzyme mechanism as a kinetic control element for designing synthetic biofuel pathways. Nature Chemical Biology 7(4), 222–227 (2011)

[6] Keasling, J., Garcia Martin, H., Lee, T.S., Mukhopadhyay, A., Singer, S.W., Sundstrom, E.: Microbial production of advanced biofuels. Nature Reviews Microbiology 19(11), 701–715 (2021)

[7] Vidal, L.S., Isalan, M., Heap, J.T., Ledesma-Amaro, R.: A primer to directed evolution: current methodologies and future directions. RSC Chemical Biology 4, 271–291 (2023)

[8] Wang, Z., Xie, D., Wu, D., Luo, X., Wang, S., Li, Y., Yang, Y., Li, W., Zheng, L.: Robust enzyme discovery and engineering with deep learning using CataPro. Nature Communications 16(1), 2736 (2025)

[9] Back, H.-m., Yun, H.-y., Kim, S.K., Kim, J.K.: Beyond the Michaelis-Menten: accurate prediction of in vivo hepatic clearance for drugs with low KM. Clinical and Translational Science 13(6), 1199–1207 (2020)

[10] Jang, H.J., Song, Y.M., Jeon, J.S., Yun, H.-y., Kim, S.K., Kim, J.K.: Optimizing enzyme inhibition analysis: precise estimation with a single inhibitor concentration. Nature Communications 16(1), 5217 (2025)

[11] Jumper, J., Evans, R., Pritzel, A., Green, T., Figurnov, M., Ronneberger, O., Tunyasuvunakool, K., Bates, R., Žídek, A., Potapenko, A., et al.: Highly accurate protein structure prediction with AlphaFold. Nature 596(7873), 583–589 (2021)

[12] Kroll, A., Ranjan, S., Engqvist, M.K., Lercher, M.J.: A general model to predict small molecule substrates of enzymes based on machine and deep learning. Nature Communications 14(1), 2787 (2023)

[13] Watson, J.L., Juergens, D., Bennett, N.R., Trippe, B.L., Yim, J., Eisenach, H.E., Ahern, W., Borst, A.J., Ragotte, R.J., Milles, L.F., et al.: De novo design of protein structure and function with RFdiffusion. Nature 620(7976), 1089–1100 (2023)

[14] Tao, H., Wang, X., Huang, S.-Y.: An interaction-derived graph learning framework for scoring protein–peptide complexes. Nature Machine Intelligence, 1–12 (2025)

[15] Heckmann, D., Lloyd, C.J., Mih, N., Ha, Y., Zielinski, D.C., Haiman, Z.B., Desouki, A.A., Lercher, M.J., Palsson, B.O.: Machine learning applied to enzyme turnover numbers reveals protein structural correlates and improves metabolic models. Nature Communications 9(1), 5252 (2018)

[16] Li, F., Yuan, L., Lu, H., Li, G., Chen, Y., Engqvist, M.K., Kerkhoven, E.J., Nielsen, J.: Deep learning-based *k*_cat_ prediction enables improved enzyme-constrained model reconstruction. Nature Catalysis 5(8), 662–672 (2022)

[17] Elnaggar, A., Heinzinger, M., Dallago, C., Rehawi, G., Wang, Y., Jones, L., Gibbs, T., Feher, T., Angerer, C., Steinegger, M., et al.: Prottrans: Toward understanding the language of life through self-supervised learning. IEEE transactions on pattern analysis and machine intelligence 44(10), 7112–7127 (2021)

[18] Rives, A., Meier, J., Sercu, T., Goyal, S., Lin, Z., Liu, J., Guo, D., Ott, M., Zitnick, C.L., Ma, J., et al.: Biological structure and function emerge from scaling unsupervised learning to 250 million protein sequences. Proceedings of the National Academy of Sciences 118(15), 2016239118 (2021)

[19] Lin, Z., Akin, H., Rao, R., Hie, B., Zhu, Z., Lu, W., Smetanin, N., Verkuil, R., Kabeli, O., Shmueli, Y., et al.: Evolutionary-scale prediction of atomic-level protein structure with a language model. Science 379(6637), 1123–1130 (2023)

[20] Honda, S., Shi, S., Ueda, H.R.: SMILES Transformer: Pre-trained molecular fingerprint for low data drug discovery. arXiv preprint arXiv:1911.04738 (2019)

[21] Edwards, C., Lai, T., Ros, K., Honke, G., Cho, K., Ji, H.: Translation between molecules and natural language. arXiv preprint arXiv:2204.11817 (2022)

[22] Yu, H., Deng, H., He, J., Keasling, J.D., Luo, X.: UniKP: a unified framework for the prediction of enzyme kinetic parameters. Nature Communications 14(1), 8211 (2023)

[23] Kroll, A., Rousset, Y., Hu, X.-P., Liebrand, N.A., Lercher, M.J.: Turnover number predictions for kinetically uncharacterized enzymes using machine and deep learning. Nature Communications 14(1), 4139 (2023)

[24] Stein, R.A.: Proteins: Structure and function. Annals of Biomedical Engineering 33, 1831–1832 (2005)

[25] Duan, C., Liu, G.-H., Du, Y., Chen, T., Zhao, Q., Jia, H., Gomes, C.P., Theodorou, E.A., Kulik, H.J.: Optimal transport for generating transition states in chemical reactions. Nature Machine Intelligence 7(4), 615–626 (2025)

[26] Boorla, V.S., Maranas, C.D.: CatPred: a comprehensive framework for deep learning in vitro enzyme kinetic parameters. Nature Communications 16(1), 2072 (2025)

[27] Satorras, V.G., Hoogeboom, E., Welling, M.: E(n) equivariant graph neural networks. In: International Conference on Machine Learning, pp. 9323–9332 (2021)

[28] Su, J., Han, C., Zhou, Y., Shan, J., Zhou, X., Yuan, F.: Saprot: Protein language modeling with structure-aware vocabulary. In: The Twelfth International Conference on Learning Representations

[29] Schramm, V.L.: Enzymatic transition states, transition-state analogs, dynamics, thermodynamics, and lifetimes. Annual review of biochemistry 80(1), 703–732 (2011)

[30] Cui, H., Su, Y., Dean, T.J., Yu, T., Zhang, Z., Peng, J., Shukla, D., Zhao, H.: Enzyme specificity prediction using cross attention graph neural networks. Nature, 1–3 (2025)

[31] Gainza, P., Wehrle, S., Van Hall-Beauvais, A., Marchand, A., Scheck, A., Harteveld, Z., Buckley, S., Ni, D., Tan, S., Sverrisson, F., et al.: De novo design of protein interactions with learned surface fingerprints. Nature 617(7959), 176–184 (2023)

[32] Ji, X., Wang, Z., Gao, Z., Zheng, H., Zhang, L., Ke, G., E, W.: Exploring molecular pretraining model at scale. In: Advances in Neural Information Processing Systems 38 (2024)

[33] Cao, D., Chen, M., Zhang, R., Wang, Z., Huang, M., Yu, J., Jiang, X., Fan, Z., Zhang, W., Zhou, H., et al.: SurfDock is a surface-informed diffusion generative model for reliable and accurate protein–ligand complex prediction. Nature Methods 22(2), 310–322 (2025)

[34] Landrum, G., Tosco, P., Kelley, B., Rodriguez, R., Cosgrove, D., Vianello, R., sriniker, Gedeck, P., Jones, G., NadineSchneider, Kawashima, E., Nealschneider, D., Dalke, A., Swain, M., Cole, B., Turk, S., Savelev, A., Vaucher, A., Wójcikowski, M., Take, I., Scalfani, V.F., Walker, R., Probst, D., Ujihara, K., Pahl, A., godin, Lehtivarjo, J., tadhurst-cdd, Bérenger, F., Bisson, J.: Rdkit/rdkit: 2024 09 1 (Q3 2024) Release. 10.5281/zenodo.13848108

[35] Chang, A., Jeske, L., Ulbrich, S., Hofmann, J., Koblitz, J., Schomburg, I., Neumann-Schaal, M., Jahn, D., Schomburg, D.: BRENDA, the ELIXIR core data resource in 2021: new developments and updates. Nucleic Acids Research 49(D1), 498–508 (2021)

[36] Wittig, U., Rey, M., Weidemann, A., Kania, R., Müller, W.: SABIO-RK: an updated resource for manually curated biochemical reaction kinetics. Nucleic Acids Research 46(D1), 656–660 (2018)

[37] Consortium, T.U.: UniProt: the universal protein knowledgebase in 2023. Nucleic Acids Research 51(D1), 523–531 (2022)

[38] Yang, K., Swanson, K., Jin, W., Coley, C., Eiden, P., Gao, H., Guzman-Perez, A., Hopper, T., Kelley, B., Mathea, M., et al.: Analyzing learned molecular representations for property prediction. Journal of Chemical Information and Modeling 59(8), 3370–3388 (2019)

[39] Bar-Even, A., Noor, E., Savir, Y., Liebermeister, W., Davidi, D., Tawfik, D.S., Milo, R.: The moderately efficient enzyme: evolutionary and physicochemical trends shaping enzyme parameters. Biochemistry 50(21), 4402–4410 (2011)

[40] Liu, Y., Hua, C., Xu, M., Zeng, T., Rao, J., Zhang, Z., Wu, R., Weng, J.-K., Coley, C.W., Zheng, S.: A geometric foundation model for enzyme retrieval with evolutionary insights. Nature Catalysis (2026)

[41] Lundberg, S.M., Lee, S.-I.: A unified approach to interpreting model predictions. Advances in neural information processing systems 30 (2017)

[42] Jendresen, C.B., Stahlhut, S.G., Li, M., Gaspar, P., Siedler, S., Förster, J., Maury, J., Borodina, I., Nielsen, A.T.: Highly active and specific tyrosine ammonia-lyases from diverse origins enable enhanced production of aromatic compounds in bacteria and saccharomyces cerevisiae. Applied and Environmental Microbiology 81(13), 4458–4476 (2015)

[43] Zhou, G., Rusnac, D.-V., Park, H., Canzani, D., Nguyen, H.M., Stewart, L., Bush, M.F., Nguyen, P.T., Wulff, H., Yarov-Yarovoy, V., et al.: An artificial intelligence accelerated virtual screening platform for drug discovery. Nature Communications 15(1), 7761 (2024)

[44] Kim, Y., Seo, P., Jeon, E., You, I., Hwang, K., Kim, N., Tse, J., Bae, J., Choi, H.-S., Hinshaw, S.M., et al.: Targeted kinase degradation via the KLHDC2 ubiquitin E3 ligase. Cell chemical biology 30(11), 1414–1420 (2023)

[45] W. Caldwell, G., Yan, Z., Lang, W., A. Masucci, J.: The IC50 concept revisited. Current topics in medicinal chemistry 12(11), 1282–1290 (2012)

[46] Haupt, L.J., Kazmi, F., Ogilvie, B.W., Buckley, D.B., Smith, B.D., Leatherman, S., Paris, B., Parkinson, O., Parkinson, A.: The reliability of estimating ki values for direct, reversible inhibition of cytochrome P450 enzymes from corresponding IC50 values: a retrospective analysis of 343 experiments. Drug Metabolism and Disposition 43(11), 1744–1750 (2015)

[47] Wei, Y., Sun, M.-m., Zhang, R.-l., Wang, L., Yang, L.-h., Shan, C.-l., Lin, J.-p.: Discovery of novel dual-target inhibitors of LSD1/EGFR for non-small cell lung cancer therapy. Acta Pharmacologica Sinica 46(4), 1030–1044 (2025)

[48] Mohammad, H.P., Smitheman, K.N., Kamat, C.D., Soong, D., Federowicz, K.E., Van Aller, G.S., Schneck, J.L., Carson, J.D., Liu, Y., Butticello, M., et al.: A DNA hypomethylation signature predicts antitumor activity of LSD1 inhibitors in SCLC. Cancer cell 28(1), 57–69 (2015)

[49] Chithrananda, S., Grand, G., Ramsundar, B.: ChemBERTa: large-scale self-supervised pretraining for molecular property prediction. arXiv preprint arXiv:2010.09885 (2020)

[50] Wang, S., Guo, Y., Wang, Y., Sun, H., Huang, J.: Smiles-bert: large scale unsupervised pre-training for molecular property prediction. In: Proceedings of the 10th ACM International Conference on Bioinformatics, Computational Biology and Health Informatics, pp. 429–436 (2019)

[51] unikei: bert-base-smiles (2025). https://huggingface.co/unikei/bert-base-smiles

[52] Wang, Y., Wang, J., Cao, Z., Barati Farimani, A.: Molecular contrastive learning of representations via graph neural networks. Nature Machine Intelligence 4(3), 279–287 (2022)

[53] Rong, Y., Bian, Y., Xu, T., Xie, W., Wei, Y., Huang, W., Huang, J.: Self-supervised graph transformer on large-scale molecular data. Advances in neural information processing systems 33, 12559–12571 (2020)

[54] Liu, S., Wang, H., Liu, W., Lasenby, J., Guo, H., Tang, J.: Pre-training molecular graph representation with 3d geometry. arXiv preprint arXiv:2110.07728 (2021)

[55] Fang, X., Liu, L., Lei, J., He, D., Zhang, S., Zhou, J., Wang, F., Wu, H., Wang, H.: Geometry-enhanced molecular representation learning for property prediction. Nature Machine Intelligence 4(2), 127–134 (2022)

[56] Zhou, G., Gao, Z., Ding, Q., Zheng, H., Xu, H., Wei, Z., Zhang, L., Ke, G.: Uni-Mol: A universal 3d molecular representation learning framework. In: The Eleventh International Conference on Learning Representations (2023)

[57] Hayes, T., Rao, R., Akin, H., Sofroniew, N.J., Oktay, D., Lin, Z., Verkuil, R., Tran, V.Q., Deaton, J., Wiggert, M., Badkundri, R., Shafkat, I., Gong, J., Derry, A., Molina, R.S., Thomas, N., Khan, Y.A., Mishra, C., Kim, C., Bartie, L.J., Nemeth, M., Hsu, P.D., Sercu, T., Candido, S., Rives, A.: Simulating 500 million years of evolution with a language model. Science 387(6736), 850–858 (2025)

[58] Lauko, A., Pellock, S.J., Sumida, K.H., Anishchenko, I., Juergens, D., Ahern, W., Jeung, J., Shida, A.F., Hunt, A., Kalvet, I., Norn, C., Humphreys, I.R., Jamieson, C., Krishna, R., Kipnis, Y., Kang, A., Brackenbrough, E., Bera, A.K., Sankaran, B., Houk, K.N., Baker, D.: Computational design of serine hydrolases. Science 388(6744), 2454 (2025)

[59] Dauparas, J., Lee, G.R., Pecoraro, R., An, L., Anishchenko, I.V., Glasscock, C.J., Baker, D.: Atomic context-conditioned protein sequence design using LigandMPNN. Nature Methods 22, 717–723 (2023)

[60] Steinegger, M., Söding, J.: MMseqs2 enables sensitive protein sequence searching for the analysis of massive data sets. Nature Biotechnology 35(11), 1026–1028 (2017)

[61] Butina, D.: Unsupervised data base clustering based on daylight’s fingerprint and tanimoto similarity: A fast and automated way to cluster small and large data sets. Journal of Chemical Information and Computer Sciences 39(4), 747–750 (1999)

[62] Tunyasuvunakool, K., Adler, J., Wu, Z., Green, T., Zielinski, M., Zídek, A., Bridgland, A., Cowie, A., Meyer, C., Laydon, A., et al.: Highly accurate protein structure prediction for the human proteome. Nature 596(7873), 590–596 (2021)

[63] Hekkelman, M.L., Vries, I., Joosten, R.P., Perrakis, A.: AlphaFill: enriching alphafold models with ligands and cofactors. Nature Methods 20(2), 205–213 (2023)

[64] Krivák, R., Hoksza, D.: P2Rank: machine learning based tool for rapid and accurate prediction of ligand binding sites from protein structure. Journal of Cheminformatics 10(1), 39 (2018)

[65] Beusekom, B., Touw, W.G., Tatineni, M., Somani, S., Rajagopal, G., Luo, J., Gilliland, G.L., Perrakis, A., Joosten, R.P.: Homology-based hydrogen bond information improves crystallographic structures in the pdb. Protein Science 27(3), 798–808 (2018)

[66] Gainza, P., Sverrisson, F., Monti, F., Rodola, E., Boscaini, D., Bronstein, M.M., Correia, B.E.: Deciphering interaction fingerprints from protein molecular surfaces using geometric deep learning. Nature Methods 17(2), 184–192 (2020)

[67] Vaswani, A., Shazeer, N., Parmar, N., Uszkoreit, J., Jones, L., Gomez, A.N., Kaiser, L, ., Polosukhin, I.: Attention is all you need. Advances in neural information processing systems 30 (2017)

[68] Kokhlikyan, N., Miglani, V., Martin, M., Wang, E., Alsallakh, B., Reynolds, J., Melnikov, A., Kliushkina, N., Araya, C., Yan, S., et al.: Captum: A unified and generic model interpretability library for pytorch. arXiv preprint arXiv:2009.07896 (2020)

